# Molecular atlas reveals the tri-sectional spinning mechanism of spider dragline silk

**DOI:** 10.1101/2022.06.21.496984

**Authors:** Wenbo Hu, Anqiang Jia, Sanyuan Ma, Guoqing Zhang, Zhaoyuan Wei, Fang Lu, Yongjiang Luo, Zhisheng Zhang, Jiahe Sun, Tianfang Yang, TingTing Xia, Qinhui Li, Ting Yao, Jiangyu Zheng, Zijie Jiang, Qingyou Xia, Yi Wang

## Abstract

We performed the first molecular atlas of natural spider dragline silk production using genome assembly for the golden orb-web spider *Trichonephila clavata* and multiomics defining for the segmented major ampullate (Ma) gland: Tail, Sac, and Duct. We uncovered a hierarchical biosynthesis of spidroins, organic acids, lipids, and chitin in the sectionalized Ma gland dedicated to fine silk constitution. The ordered secretion of spidroins was achieved by the synergetic regulation of epigenetic and ceRNA signatures for genomic group-distributed spidroin genes. Single-cellular and spatial RNA profiling identified ten cell types with partitioned functional division determining the tri-sectional organization of the Ma gland. Convergent evolution and genetic manipulation analyses further validated that this tri-sectional architecture of the silk gland was analogous in silk-spinning animals and inextricably linked with silk formation. Our study provided multiple levels of data that significantly expand the knowledge of spider dragline silk generation and may eventually benefit spider-inspired fiber innovations.

## Introduction

Spiders are abundant generalist arthropod predators with more than fifty thousand extant species^1, 2^. Spider silks are natural high-performance proteinaceous fibers and are crucial to spider survival and reproduction^3, 4^. A fascinating spider-salient trait of particular economic interest is silk production, mainly due to the exceptional properties, including high tensile strength and toughness, low density, self-powered rotational actuation, and biocompatibility, of these fibers^5–7^. Numerous works have persistently tried to emulate natural spidroin production and spinning processes for the biomimetic generation of artificial materials with spider silk-like properties^8–11^. Thus, an in-depth understanding of the molecular mechanisms acting in the natural silk production system is valuable for unraveling the mystery of spider silk.

The major ampullate (Ma) gland of the orb-web spider is a model system in silk production research due to the relatively large size and especially the impressive properties of its product, dragline silk^12^. It was observed that the Ma gland can be divided into three macroscopic parts, the Tail, the Sac, and the Duct, with gradient pH values, ion concentrations, and shear forces^13–15^, in which the liquid silk protein was synthesized and stored at a very high concentration in the Tail and Sac, then transformed into insoluble fiber through the Duct^14, 16^. In line with this, most attempts at dragline silk-inspired fiber innovations have largely employed the silk proteins and microenvironment produced by the Ma gland^13–16^. For example, of the major ampullate spidroins (MaSps), recombinant repetitive motifs were constructed to achieve specific physical properties, including strength, extensibility, and stickiness, for silk^17–19^. Spider silk-constituting elements (SpiCEs), nonspidroin proteins, are utilized to increase tensile strength in the case of composite silk films^20–22^. In addition, the microfluidics device designed to realize the closest equivalent to natural ionic and pH conditions allows the fibers to be directly pulled out from the outlet and then reeled in air, as in native spinning^23–25^. To enable the future reproduction of dragline silk, we need to gain more details of the Ma gland, the cellular architecture and molecular function, as well as the biocomposition and formation process of dragline silk^26–28^.

Thus far, comprehensive genomic analyses of orb-web spiders have parsed the molecular traits of MaSps as well as other nonspidroin proteins composing dragline silk^17, 21, 28–31^. Nevertheless, genetically mediated Ma gland mechanisms underlying dragline silk production remain poorly understood. Herein, we present a chromosome-scale reference genome for the golden orb-web spider *Trichonephila clavata.* This high-quality genome combined with multiomics approaches, including RNA-seq, liquid chromatography–mass spectrometry (LC–MS), ATAC-seq, bisulfite-seq, whole-transcriptomics (WT), single-cell (SC) RNA-seq, and spatial transcriptomics (ST), generate the first detailed anatomic definitions for the segmented Ma gland (the Tail, Sac, and Duct) based exclusively on segment-specific and dragline silk-related gene expression classification, which allows us to entirely delineate the molecular atlas of silk production processes within the Ma gland, thereby visualizing a tri-sectional generation mechanism of dragline silk. These multiomics datasets are accessible in SpiderDB (https://spider.bioinfotoolkits.net) and valuable for future explorations of the evolutionary origins of silk production strategies and the creation of biomimetic spider silks.

## Results

### Chromosomal-scale genome assembly and full spidroin gene set of *T. clavata*

The golden orb-web spider, *T. clavata* exhibited extreme sexual size dimorphism and constructed a large and impressive orb web (Fig. 1a). To assemble a high-quality genome, we first performed cytogenetic analysis of *T. clavata* and found 2n = 26 in females and 2n = 24 in males, comprising eleven pairs of autosomal elements and unpaired sex chromosomes (X_1_X_1_X_2_X_2_ in females and X_1_X_2_ in males) (Fig. 1a). Then, DNA from adult female *T. clavata* was used to generate long-read (Oxford Nanopore Technologies (ONT)), short-read (Illumina), and Hi-C data. Our sequential assembly approach (Supplementary Fig. 1c) resulted in a 2.63 Gb genome with a scaffold N50 of 202.09 Mb and a Benchmarking Universal Single-Copy Ortholog (BUSCO) genome completeness score of 93.70% (Table 1). Finally, the genome was assembled into 13 pseudochromosomes. Sex-specific Pool-Seq analysis from male and female spiders indicated that Chr12 and Chr13 were sex chromosomes (Fig. 1b; Supplementary Fig. 2). Based on the MAKER2 pipeline^32^ (Supplementary Fig. 1e), we annotated 37,607 protein-encoding gene models and predicted repetitive elements with a collective length of 1.42 Gb, accounting for 53.94% of the genome.

**Figure 1.**
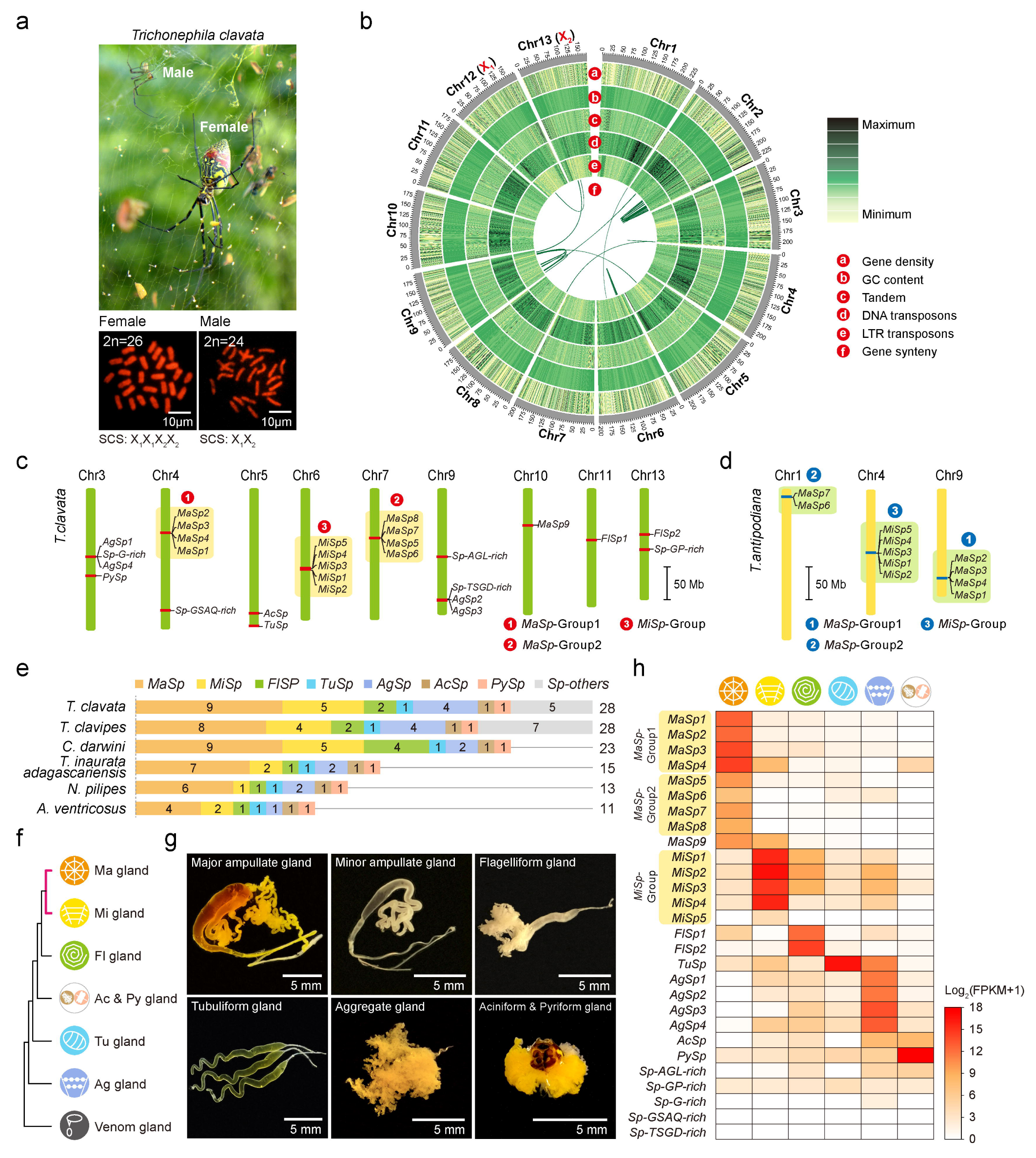
Chromosomal-scale genome assembly and full spidroin gene set of *T. clavata*. **a**, Photograph of *T. clavata* showing an adult female and an adult male on the golden orb-web (above) and the karyotypes of female and male (below). SCS represents the sex chromosome system. **b**, Circular diagram depicting the genomic landscape of the 13 pseudochromosomes (Chr1*–*13 on an Mb scale). **c**, Twenty-eight *T. clavata* spidroin genes anchored on chromosomes. **d**, Spidroin gene groups in *T. antipodiana*. **e**, Spidroin gene catalog of six orb-web spider species. **f**, Expression clustering of silk glands (major and minor ampullate- (Ma and Mi), flagelliform- (Fl), tubuliform- (Tu), aggregate- (Ag), and aciniform & pyriform (Ac & Py) glands) and venom glands. The pink line shows the closest relationship between Ma and Mi glands. **g**, Morphology of *T. clavata* silk glands. **h**, Expression patterns of 28 spidroin genes in different types of silk glands.

**Table 1.** Characteristics of the T. clavata genome assembly.

|  |  |
| --- | --- |
| **Estimated genome size (bp)** | **2,722,533,452** |
| Heterozygosity (%) | 1.44 |
| **Length of genome assemble (bp)** | **2,730,479,360** |
| **Total length of scaffolds (bp)** | **2,732,859,360** |
| N50 of scaffolds (bp) | 202,090,426 |
| N90 of scaffolds (bp) | 173,619,284 |
| **Total length of contigs (bp)** | **2,730,479,360** |
| N50 of contigs (bp) | 1,788,282 |
| N90 of contigs (bp) | 215,983 |
| GC content (%) | 31.80 |
| Complete BUSCOs | 93.70 |
| **Total length of pseudomolecules (bp)** | **2,629,098,020** |
| Number of genes | 37,607 |
| Repeat content (%) | 53.94 |

To identify *T. clavata* spidroin genes, we searched the annotated gene models for sequences similar to 443 published spidroins (Supplementary Table 6) and performed phylogenetic analysis of the putative spidroin sequences for classification (Supplementary Fig. 12a). Based on the principle that a typical spidroin gene consists of a long repeat domain sandwiched between the nonrepetitive N/C-terminal domains^27^, these candidates were further validated and reconstructed using full-length transcript isoform sequencing (Iso-seq) and transcriptome sequencing (RNA-seq) data. We then identified 28 spidroin genes, among which 15 were full length. This full set of spidroin genes was located on nine of 13 *T. clavata* chromosomes. Interestingly, we found that the *MaSp1–4*, *MaSp5–8*, and *MiSp1–5* genes were distributed in three independent groups (Fig. 1c). Of note, such distribution was also detected in the chromosomal-level *T. antipodiana* genome^31^ (Fig. 1d), supporting the reliable grouping results in our study. When comparing the spidroin gene catalog of *T. clavata* and five other sequenced orb-web spider species^21, 30, 33, 34^ (Fig. 1e), we found that *T. clavata* and *T. clavipe* possessed the same and largest number of spidroin genes.

**Figure 2.**
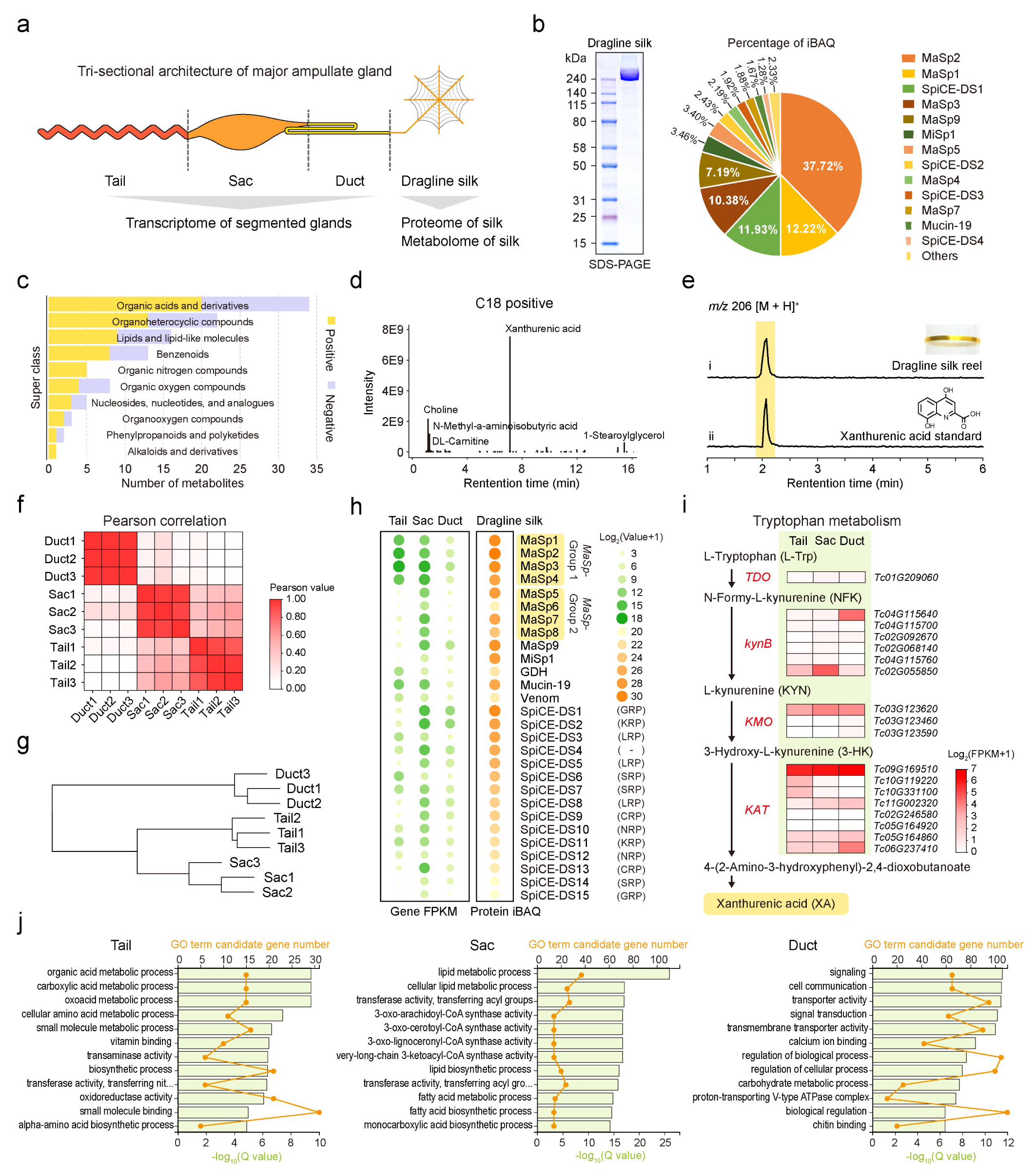
Dragline silk origin and the functional character of the tri-section Ma gland. **a**, Schematic image of Ma gland segmentation. **b**, Sodium dodecyl sulfate–polyacrylamide gel electrophoresis (SDS–PAGE) (left) and LC–MS (right) analyses of dragline silk protein. **c**, LC–MS analyses of the metabolites in dragline silk. **d**, Classification of the identified metabolites. **e**, LC–MS analyses of the golden extract from *T. clavata* dragline silk. The extracted ion chromatograms (EICs) were extracted at *m/z* 206 [M + H]^+^ for xanthurenic acid. **f**, Pearson correlation of different Ma gland segments (Tail, Sac, and Duct). **g**, Expression clustering of the Tail, Sac, and Duct. **h**, Combinational analysis of the transcriptome and proteome showing the expression profiling of the dragline silk gene in the Tail, Sac, and Duct. **i**, Concise biosynthetic pathway of xanthurenic acid (tryptophan metabolism) in the *T. clavata* Ma gland. Gene expression levels mapped to tryptophan metabolism are shown at three segments of the Ma gland. Enzymes involved in the pathway are marked in red, and the genes encoding the enzymes are shown beside them. **j**, Gene Ontology (GO) enrichment analysis of Ma gland segment-specific genes showing the biological functions of Tail, Sac, and Duct. The top 12 significantly enriched GO terms were exhibited for each segment of the Ma gland.

Next, all morphologically distinct glands (major and minor ampullate- (Ma and Mi), flagelliform- (Fl), tubuliform- (Tu), and aggregate (Ag) glands) were cleanly and separately dissected from adult female *T. clavata* except for the aciniform and pyriform glands (Fig. 1g), which because of their proximal anatomical locations could not be cleanly separated and were therefore treated as a combined sample (aciniform & pyriform gland (Ac & Py)). We performed whole-gene expression clustering analysis of all silk glands and found that the Ma and Mi glands showed the closest relationship in morphological structure and gene expression (Fig. 1f,h). We noted that the expression profiles of spidroin genes largely agreed with their putative ascribed roles in their corresponding morphologically distinct silk glands, such as *MaSps* in the Ma gland (Fig. 1h). Meanwhile, some spidroin transcripts were expressed in several silk glands, such as *MiSps* and *TuSp* (Fig. 1h). The unclassified spidroin genes did not appear to be gland-specific in expression, such as Sp-GP-rich (Fig. 1h). These observations provide clues regarding the function of spidroin genes in different silk glands.

In summary, the chromosomal-scale genome of *T. clavata* allowed us to obtain detailed structural and location information of whole-spidroin genes. We also found a relatively diverse set of spidroin genes and a group-distributed feature of *MaSps* and *MiSps* in *T. clavata*.

### Dragline silk origin and the functional character of the tri-section Ma gland

We performed integrated analyses of the dragline silk proteome, dragline silk metabolome, and Ma gland transcriptome to characterize the molecular details of the tri-section Ma gland in the dragline silk secreting process (Fig. 2a). Sodium dodecyl sulfate–polyacrylamide gel electrophoresis (SDS–PAGE) analysis of *T. clavata* dragline silk showed a thick band above 240 kDa, implying a smaller variety of total proteins (Fig. 2b). Subsequent LC–MS analysis identified ten spidroins and 18 nonspidroin proteins from this silk, containing nine MaSps, a MiSp, a glucose dehydrogenase (GDH), a mucin-19, a venom, and 15 SpiCEs of dragline silk (SpiCE-DS) with unknown functional annotation. Among these proteins, we found that the core protein components of dragline silk were MaSp2 (37.7%), MaSp1 (12.2%), SpiCE-DS1 (11.9%, also known as SpiCE-NMa1 in a previous study^21^), MaSp3 (10.4%), and MaSp9 (7.2%) in order of the percentage of iBAQ, accounting for approximately 80% of the total protein abundance of dragline silk (Fig. 2b). This result provided potential protein components that might be highly correlated with excellent strength and toughness.

We then identified a total of 180 metabolic components from *T. clavata* dragline silk (Supplementary Table 12). Based on certain features of metabolites, they were classified into ten categories: organic acid, organoheterocyclic compound, lipid, benzenoid, organic nitrogen, organic oxygen, nucleoside, organooxygen, phenylpropanoid and polyketide, and alkaloid (Fig. 2c; Supplementary Table 13). The golden color is the most obvious feature of *T. clavata* dragline silk. We noted that xanthurenic acid (XA, a yellow pigment) was most abundant (Fig. 2d), while other yellow pigments (such as carotenoids and flavonoids) were not detected in our analysis, implying that XA was the major pigment of golden silk. Moreover, the XA was further confirmed by LC–MS analysis of the golden extract (Fig. 2e), consistent with a recent report^21^.

To explore the origin of dragline silk components from the tri-section Ma gland, we focused on the transcriptomic features of the Tail, Sac, and Duct. We determined that the gene expression profiles of the Tail and Sac were more correlated than those of the Duct (Fig. 2f,g; Supplementary Fig. 13c), implying a similar molecular function of the Tail and Sac. Further transcriptomic and proteomic combination analyses showed the expression pattern of 28 dragline silk protein transcripts in the Tail, Sac, and Duct (Fig. 2h; Supplementary Table 10). As shown in Fig. 2h, we could accurately determine where dragline silk proteins from. It is worth noting that *MaSp1–4* (*MaSp*-Group1) were highly coexpressed in the Tail and Sac, and *MaSp5–8* (*MaSp*-Group2) were highly coexpressed only in the Sac, indicating that the Tail and Sac were major silk-secreting segments. We then used the tri-section Ma gland datasets to trace the source of metabolite XA. We found that the genes encoding key enzymes involved in the XA biosynthesis (tryptophan metabolism) pathway were activated in all three Ma gland segments (Fig. 2i), especially a kynurenine aminotransferase gene (*KAT*, *Tc09G169510*) that encoded the primary enzymes catalyzing the transamination of 3-hydroxy-L-kynurenine (3-HK) to XA, suggesting that XA was secreted by the Tail, Sac, and Duct.

**Figure 3.**
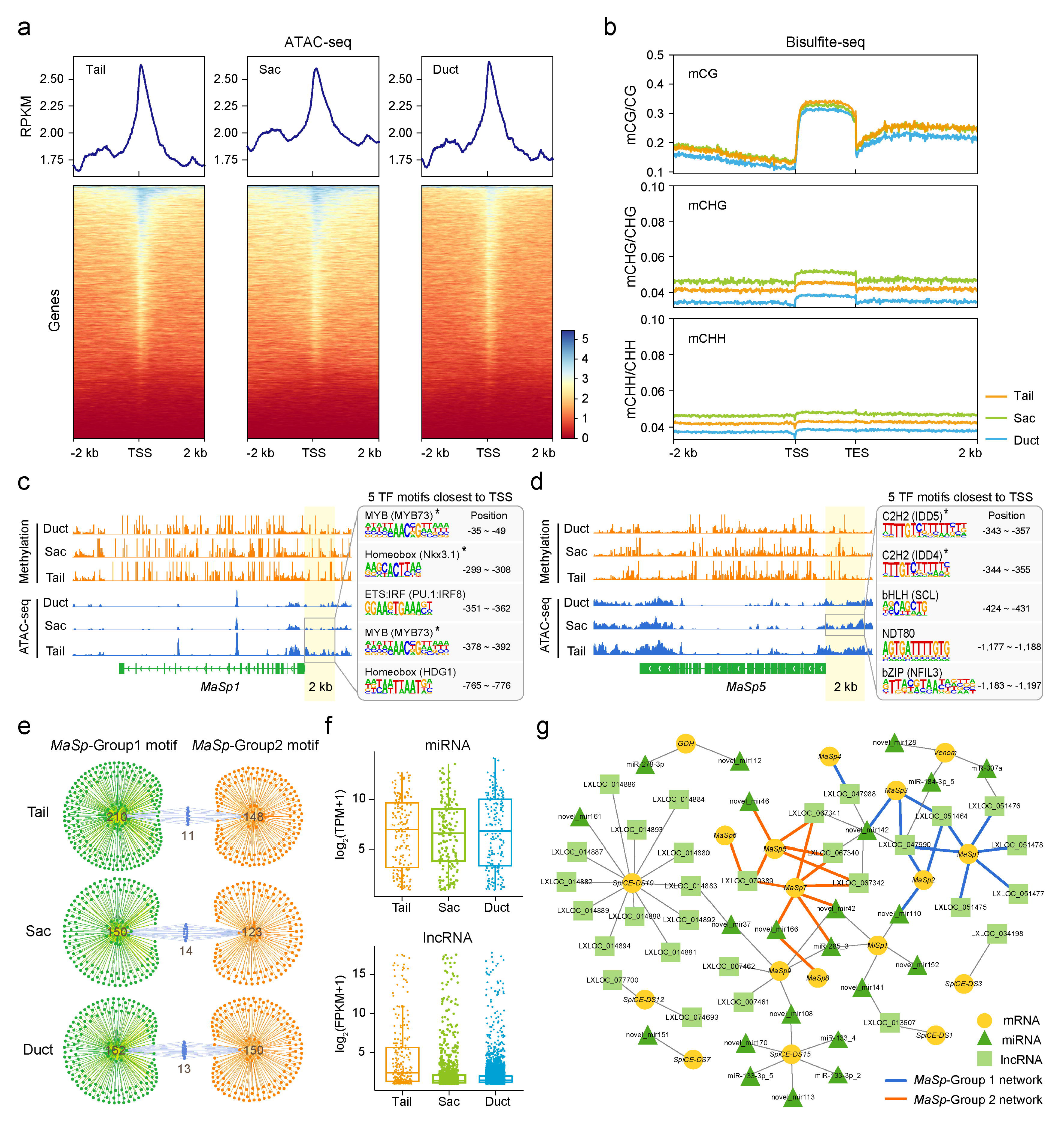
Comprehensive epigenetic features and ceRNA network of the tri-section Ma gland. **a**, Metagene plot of ATAC-seq signal and heatmap of the ATAC-seq read densities in Tail, Sac, and Duct. **b**, Metagene plot of DNA methylation levels for CG/CHG/CHH contexts in Tail, Sac, and Duct. **c**, **d**, Screenshot of the *MaSp1*(**c**) and *MaSp5* (**d**) genes with methylation and ATAC-seq tracks in Tail, Sac, and Duct. The enriched TF motif for the indicated peak set (2 kb upstream of the TSS) is listed to the right and sorted by position. Asterisks represent the coenriched TF motif within the corresponding *MaSp* group. **e**, Venn network of enriched TF motifs between *MaSp*-Group1 and *MaSp*-Group2. **f**, The expression levels of miRNAs and lncRNAs in the Tail, Sac, and Duct. **g**, Construction of the ceRNA network of the dragline silk gene.

Next, we characterized the specific biological functions of the Tail, Sac, and Duct related to dragline silk production. The Gene Ontology (GO) terms were assigned to classify the functions of Ma gland segment-specific genes (Supplementary Table 14). We found that the significantly enriched GO terms in the Tail were mainly related to the synthesis of organic acids (the largest group in the silk metabolome), in the Sac were mainly related to the synthesis of lipids (the third-largest group in the silk metabolome), and in the Duct were related to ion (Ca^2+^ and H^+^) exchange and chitin synthesis (Fig. 2j).

Taken together, our results demonstrate a tri-sectional generation process of dragline silk in the Ma gland. We established a genetic relationship between dragline silk components and the Tail, Sac, and Duct glands.

### Comprehensive epigenetic features and ceRNA network of the Ma gland tri-section

Based on the Ma gland RNA-seq data, we found that the transcription of dragline silk genes was incredibly efficient in the Tail and Sac, and the total FPKM of these genes accounted for 47.49% and 34.33% of whole genes, respectively; however, in the Duct, they only accounted for 0.76% (Supplementary Table 11). In particular, the *MaSps* within the group were highly coexpressed in the specific segment of the Ma gland (Fig. 2h). To better understand the transcriptional regulatory mode of these genes, we first investigated genome-wide chromatin accessibility using ATAC-seq in the Tail, Sac, and Duct. We found that the Tail (mean RPKM: 1.78) and Sac (mean RPKM: 2.04) plots showed genes with more accessible chromatin than the Duct (mean RPKM: 1.59) plots (Fig. 3a), suggesting a potential regulatory role of chromatin accessibility in the transcription of dragline silk genes. We then analyzed the genome-wide DNA methylation level in the Tail, Sac, and Duct. We found the most DNA methylation in the CG context (beta value: 0.12 in Tail, 0.13 in Sac, and 0.10 in Duct) and only a small amount in the CHH (beta value: 0.04 in Tail, 0.05 in Sac, and 0.03 in Duct) and CHG (beta value: 0.04 in Tail, 0.05 in Sac, and 0.04 in Duct) contexts (Fig. 3b). Overall, there was no significant difference in methylation levels among the Tail, Sac, and Duct.

Next, visualizing the ATAC-seq and methylation datasets for two *MaSp* groups in a genome browser revealed a reverse trend of peak signals (Fig. 3c,d; Supplementary Fig. 16). We analyzed the enriched TF motif for the ATAC-seq peak set 2 kb upstream of the transcriptional start site (TSS) (Supplementary Table 15). We identified nine enriched Tail- and Sac-specific TF motifs for *MaSp1* (within *MaSp*-Group 1) and 13 enriched Sac-specific TF motifs for *MaSp5* (within *MaSp*-Group 2) (Supplementary Fig. 17a,b). Interestingly, we noted that the enriched TF motifs closest to TSS, for example, MYB and homeobox motifs for *MaSp*-Group 1 and two C2H2 motifs for *MaSp*-Group 1, were shared within the *MaSp*-Group (Fig. 3c,d). However, the Venn network enriched TF motifs between *MaSp*-Group1 and *MaSp*-Group2 showed little commonality in the variety of Tail, Sac, and Duct (Fig. 3e). In that sense, we concluded a common regulatory pattern within each *MaSp* group but a differentiated regulatory pattern among the two *MaSp* groups.

To investigate the impact of ceRNAs corresponding to the regulation of dragline silk genes, we identified a total of 527 miRNAs (179 in Tail, 167 in Sac, 181 in Duct) and 10,110 lncRNAs (240 in Tail, 982 in Sac, and 4,808 in Duct) by preforming the WT of the tri-section Ma gland (Fig. 3f). From these data, we determined the potential lncRNA–miRNA–mRNA interaction pairs by using miRanda and RNAhybrid algorithms. Then, we visualized the ceRNA network of dragline silk genes, which consisted of 28 lncRNAs, 21 miRNAs, and 13 mRNAs (Fig. 3g). Remarkably, we noted that the ceRNA networks of *MaSp1–4* were tightly clustered, as was *MaSp5–8*; in addition, the ceRNA networks of the two *MaSp* groups were independent of each other (Fig. 3g). These results further revealed the specific coregulatory pattern of *MaSp*-group genes in the Ma gland.

In summary, we detected an abundance of epigenetic and ceRNA signatures associated with the efficient and segment-specific transcription of dragline silk genes. Our data suggested a differential regulation strategy of the tri-section Ma gland dedicated to achieving the hierarchical gene expression of the two *MaSp* groups.

### Single-cell spatial architecture at the whole-Ma gland scale

To further explore the cytological basis related to the hierarchical organization of the Ma gland, we generated a whole-Ma gland molecular atlas by capturing the single-cellular and spatial patterns of gene expression in the *T. clavata* Ma gland using scRNA-seq and ST. We first identified cellular profiles of the Ma gland by performing clustering analyses of a resource of 9,349 high-quality cells generated from scRNA-seq on dissociated fresh tissue. Using UMAP clustering, we split the captured single cells into ten compartments (Fig. 4a). Based on the GO analysis of cluster-specific marker genes combined with the expression profiles of cluster- and segment-specific genes in each cell type (Supplementary Fig. 21 and Supplementary Note “Cell type annotation”), we carried out the fine annotations of ten SC clusters, namely, cluster 1: Ma gland origin cell (MaGO), cluster 2: MaSp-Group synthesis cell (MG1S), cluster 3: Chitin synthesis cell (CS), cluster 4: Unknown cell, cluster 5: Ampullate lumen skeleton cell (ALS), cluster 6: Ion transport cell (IT), cluster 7: Lipid synthesis cell (LS), cluster 8: pH adjustment cell (PA), cluster 9: MaSp-Group 2 synthesis cell I (MG2S I), and cluster 10: MaSp-Group 2 synthesis cell II (MG2S II), and delineated their sources from the Tail (clusters 1 and 2), Sac (clusters 1, 2, 5, 7, 9, 10), and Duct (clusters 1, 3, 4, 6, 8) (Fig. 4a).

**Figure 4.**
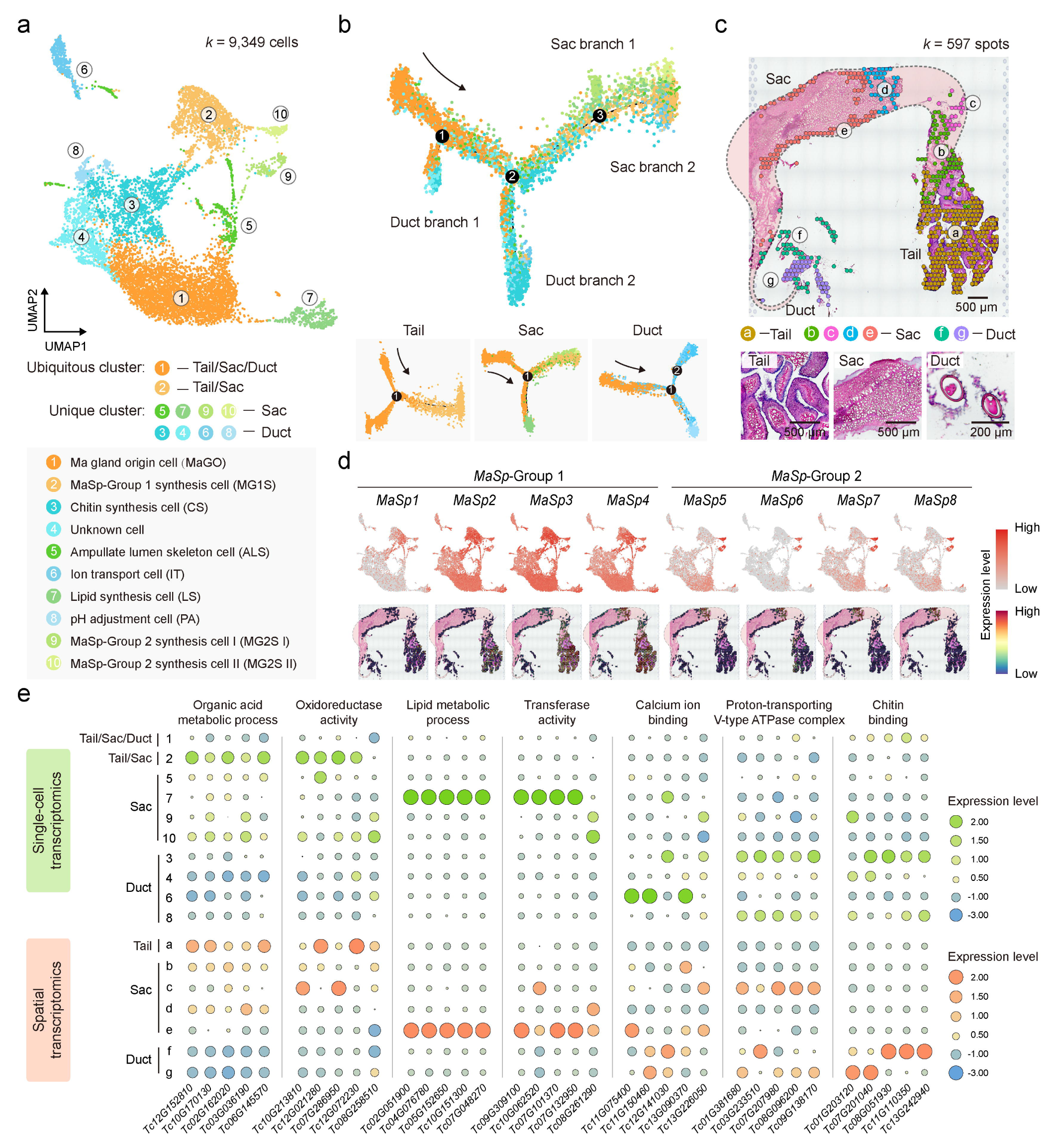
Single-cell spatial architecture at the whole-Ma gland scale. **a**, UMAP of cell types of the Ma gland and their grouping into ten cell clusters. The ubiquitous clusters are shown in yellow series, the Sac clusters in green series, and the Duct clusters in blue series. **b**, Pseudotime trajectory of all 9,349 Ma gland cells. Each dot indicates a single cell, color-coded by the cluster as in (a). The black arrow indicates the start of the trajectory. **c**, Hematoxylin and eosin (H.E.) staining of Ma gland sections and unbiased clustering of spatial transcriptomic (ST) spots. Dotted lines depict the outline of the Ma gland. **d**, UMAP and ST feature plots of expression for genes in the *MaSp* group. **e**, Heatmap showing the expression of Ma gland segment-specific genes in each cell type and each ST cluster.

To discern how Tail, Sac, and Duct arise in the Ma gland, we ordered all single cells according to pseudotime and constructed a developmental trajectory. This resulted in a continuum of cells with three distinct branch points. We found that a set of cells from cluster 1 (ubiquitous SC cluster) assembled at the beginning of pseudotime and gradually bifurcated into four ends representing two segments (the Sac and Duct) of the Ma gland, with the clusters arranged at different branch sites (Fig. 4b). We further investigated the developmental trajectory of the Tail, Sac, and Duct separately. As expected, most cells from cluster 1 assembled at the beginning of pseudotime, while the Sac cells (clusters 2, 5, 7, 9, and 10) and the Duct cells (clusters 3, 4, 6, 8) grouped into different branches (Fig. 4b; Supplementary Fig. 23). These results provide insights into the differentiation trajectory of Ma gland cells during cell state transitions.

We next assessed the spatial organization of cell populations in the Ma gland section. This dataset contained gene expression information across a resource of 597 spots (Fig. 4c). After analyzing the transcriptional signatures of ST spots, we identified seven spot clusters (one in the Tail, four in the Sac, and two in the Duct). As a first demonstration of the single-cellular and spatial expression patterns for whole genes in the Ma gland, we visualized the expression of the *MaSp*-Group1 and *MaSp*-Group2 genes in UMAP and ST feature plots to better locate silk protein secretion cells within our captured cell populations. Interestingly, we found that the *MaSp*-Group1 genes were prominently expressed in the Tail and Sac clusters (SC clusters 1, 2, 5, 7, 9, and 10 and ST cluster “a–f”), and the *MaSp*-Group2 genes were predominantly expressed in the Sac clusters (SC cluster 9–10 and ST cluster “d”) (Fig. 4d; Supplementary Fig. 22a). These genes exhibited cellular and spatial cluster-specific patterns in scRNA-seq and ST, consistent with the expression and regulatory results (Fig. 2h, 3c,d, and 3g).

To characterize the identified SC and ST clusters related to the molecular function of the Ma gland, we selected 35 well-established segment-specific genes to perform expression analysis for scRNA-seq and ST datasets. We found that SC cluster 2 and ST cluster “a” were major sets with the function of organic acid metabolic process and oxidoreductase activity, SC cluster 7 and ST cluster “e” were major sets with the function of lipid metabolic process and transferase activity, SC cluster 6 and ST cluster “f” were major sets with the function of calcium ion binding, SC cluster 3/8 and ST cluster “c/g” were major sets with the function of proton-transporting V-type ATPase complex, and SC cluster 3 and ST cluster “f” were major sets with the function of chitin binding (Fig. 4e; Supplementary Fig. 25 and 26).

**Figure 5.**
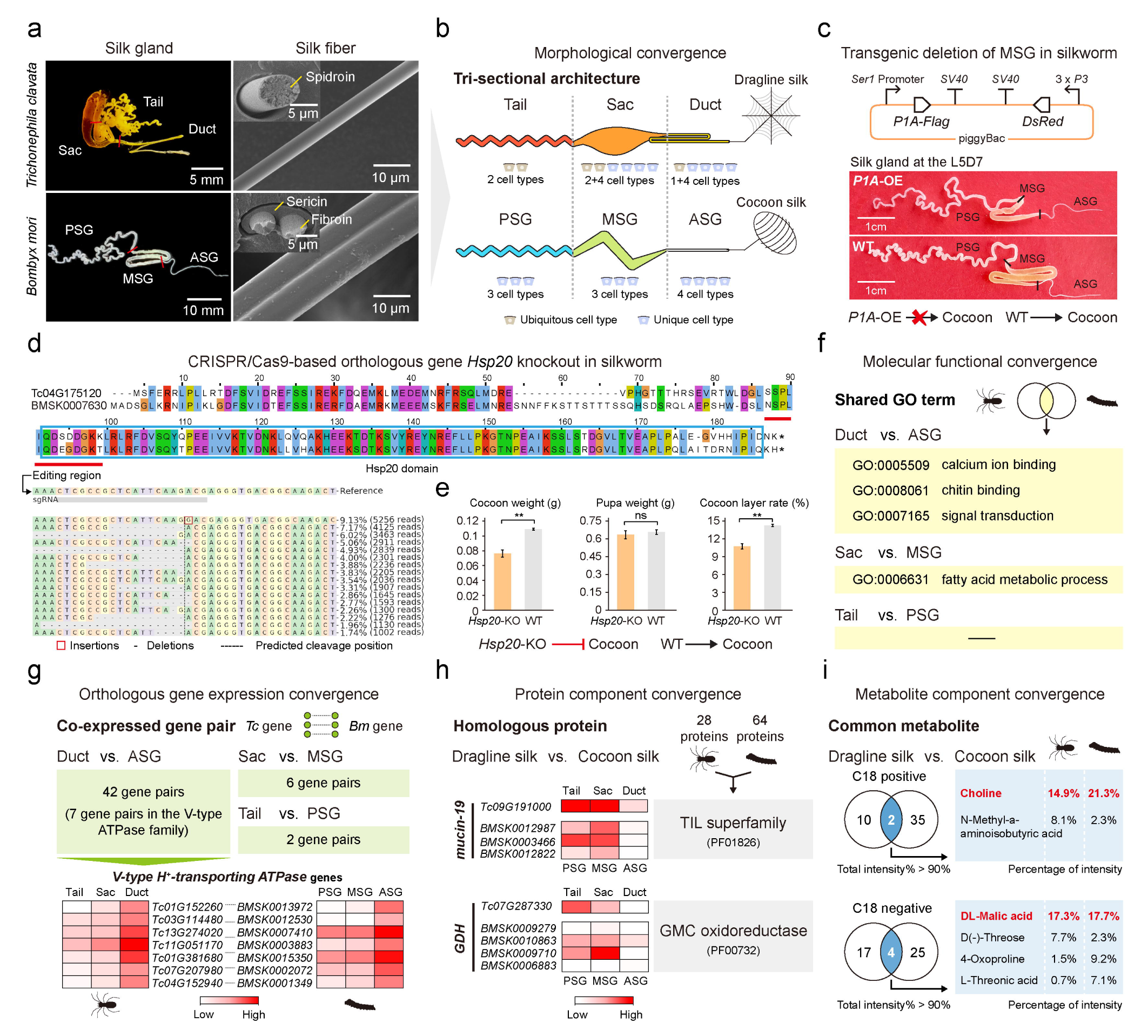
Convergent evolution of the tri-section silk gland between *T. clavata* and *B. mori*. **a**, Macroscopic appearance of the *T. clavata* Ma gland, *T. clavata* dragline silk, *B. mori* silk gland, and *B. mori* silk. **b**, Schematic illustration showing morphological convergence of the *T. clavata* Ma gland with Tail, Sac, and Duct indicated (above) and the *B. mori* silk gland with posterior silk gland (PSG), middle silk gland (MSG), and anterior silk gland (ASG) indicated (below). **c**, Construction of the piggyBac transgenic vector (above) and silk gland phenotype of the *P1A*-OE strain in silkworm (below). L5D7 represents the 7th day of the fifth instar. The arrow indicates the silk production process. Red-cross indicates that the process was blocked. **d**, Sequence alignment of the orthologous protein Hsp20 (Tc04G175120 and BMSK0007630) in spider and silkworm. The red line indicates the editing region. The arrow shows the distribution of identified alleles around the cleavage site for the sgRNA. **e**, Cocoon weight, pupa weight, and cocoon layer rate performance of the CRISPR/Cas9-based *Hsp20* knockout strain in silkworm. The arrow indicates the silk production process. The red line ending in a crossbar indicates that the process was suppressed. **f**, Molecular functional convergence of the *T. clavata* Ma gland and the *B. mori* silk gland. **g**, Orthologous gene expression convergence of the *T. clavata* Ma gland and the *B. mori* silk gland. **h**, Protein component convergence of silk produced by *T. clavata* and *B. mori*. **i**, Metabolite component convergence of silk produced by *T. clavata* and *B. mori*. The metabolites with a total intensity percentage of more than 90% were analyzed. The major metabolites in *T. clavata* dragline silk and *B. mori* cocoon silk are marked with red.

In summary, we generated a detailed anatomic definition of the Ma gland. These results revealed a simple cell type within the Tail and multiple cell types within the Sac and Duct, highlighting the developmental and functional differentiation of the Ma gland in the tri-section.

Convergent evolution of the tri-section silk gland between *T. clavata* and *B. mori*

The spider and silkworm have distinct phylogenetic positions in the phylum Arthropoda, but both evolved specialized glands for silk-spinning^35^. We noted a morphological convergence of the silk-spinning gland between the *T. clavata* Ma gland (Tail, Sac, and Duct) and the *B. mori* silk gland (posterior silk gland (PSG), middle silk gland (MSG), and anterior silk gland (ASG)), revealing a one-to-one corresponding tri-sectional architecture (Fig. 5a,b). We also found a similar number of silk gland cell types between *T. clavata* and *B. mori*^36^ but differentiated annotations except for the chitin-related process in the Duct/ASG (Fig. 5b; Supplementary Fig. 29). Our previous studies reported causal effects of structural deficiency of the PSG and ASG on silkworm spinning defects^37, 38^. Since the genetic manipulation system for spiders has not yet been established, we used *ser1* promoter-driven transgenic overexpression (OE) of a butterfly cytotoxin (pierisin-1A, P1A^39^) to generate an MSG-deficient silkworm (Fig. 5c). We found that the MSG of the *P1A*-OE strain was successfully truncated and that the *P1A*-OE strain failed to spin a cocoon, consistent with the phenotype of PSG- and ASG-deficient silkworms. Our results further demonstrated that the tri-sectional architecture of the silk gland was essential to silk spinning.

Using whole-genome blastp alignment, we then generated 9,593 and 7,355 orthologous genes between *T. clavata* and *B. mori*. From these assigned gene ortholog pairs, we selected an *Hsp20* pair (*Tc04G175120* and *BMSK0007630*) with a high sequence identity of 87.1% (Fig. 5d), which encodes a small heat shock protein that acts as a protein chaperone to protect other proteins against misfolding and aggregation^40^. We noted that *Hsp20* was expressed in the silk glands of both *T. clavata* and *B. mori* and simultaneously was the marker gene of SC cluster 5 of the *T. clavata* Ma gland (Supplementary Table 17). We then performed CRISPR/Cas9-based knockout (KO) of *Hsp20* in silkworm and successfully detected indels at the target site of *Hsp20* sgRNA (Fig. 5d). Interestingly, we found that the silk production (cocoon weight and cocoon layer rate) of the *Hsp20*-KO strain was significantly decreased (Fig. 5e). From these results, we concluded that the orthologous *Hsp20* played a positive role in silk production in both spider and silkworm.

To further investigate the molecular signatures of silk spinning for convergent evolution in both spider and silkworm, we next performed comparative transcriptome, proteome, and metabolome analyses of tri-section silk glands and silk between *T. clavata* and *B. mori*. The shared GO terms were identified in each corresponding silk gland region of *T. clavata* and *B. mori*. We found that three GO terms (calcium ion binding, chitin binding, and signal transduction) were commonly enriched in the Duct/ASG, and one GO term (fatty acid metabolic process) was commonly enriched in the Sac/MSG (Fig. 5f). Using a customized pipeline (Supplementary Fig. 28a), we found that 42, 6, and 2 pairs of orthologous genes were coexpressed in the Duct/ASG, Sac/MSG, and Tail/PSG, respectively (Fig. 5g; Supplementary Table 20). Interestingly, we found that seven gene ortholog pairs involved in the *V-type ATPase* family, encoding key factors in regulating silk fibrillogenesis^41^, were significantly upregulated in both the Duct and ASG (Fig. 5g). Our results indicated a higher consistency of the Duct/ASG in molecular functions compared with the Sac/MSG and Tail/PSG.

We then explored whether the components of spider and silkworm silks were convergent by performing Venn analyses of the silk proteome and metabolome across the two species. We found the coexistence of mucin-19 and GDH in *T. clavata* dragline silk and *B. mori* cocoon silk, and more interestingly, these proteins were mainly synthesized and secreted by Tail/PSG and Sac/MSG (Fig. 5h). We also identified six common metabolites in these two silks, including choline, N-methyl-a-aminoisobutyric acid, DL-malic acid, D(-)-threose, 4-oxoproline, and L-threonic acid (Fig. 5i). Among them, choline, a component of phospholipids in cell membranes, and DL-malic acid, a preservative or pH-adjuster, were both major metabolites of dragline silk and cocoon silk (Fig. 5i). We therefore speculated that silk secretion from gland cells to the lumen was accompanied by choline release from the cell membrane, and in nature, silk anti-rot/-bacteria was realized by DL-malic acid.

In summary, we established a comprehensive molecular comparison of the tri-section silk gland between spider and silkworm. The shared characteristics represented convergent evolution under similar selective pressures in silk spinning and yielded rich insights into silk gland function and the silk biosynthesis process.

## Discussion

Much of our current understanding of spider silk formation is based on physical and material studies that have provided only a partial picture of its nature^42, 43^. A comprehensive molecular biological atlas of the Ma gland and dragline silk is fundamental to uncovering how spiders make dragline silk fibers^11, 44^. Additionally, such a detailed basis of silk formation will be valuable for ultratough fiber innovation^5, 8, 42, 45^. Herein, we report the first chromosome-scale reference genome assembly of *T. clavata*, with a final assembly length of 2.63 Gb. This high-quality genome combined with multiomics analyses enabled visualization of silk biosynthesis processes in the tri-sectional Ma gland.

We found genomic clues that contributed to the hierarchically ordered biosynthesis of spidroins. Spidroins are core components of spider silks, the relative abundance of which is thought to exert an important influence on the mechanical properties of the corresponding fibers^27, 46, 47^. We document a novel characteristic of spidroin genes in the *T. clavata* genomes: the *MaSp1–4*, *MaSp5–8*, and *MiSp1–5* genes are distributed in three distinct groups. In addition, we further demonstrated that the *MaSps* within the same group exhibited a concert SC and ST expression profile in the tri-section Ma gland and a coregulatory pattern at the epigenetic level and ceRNA level. As more high-quality spider genomes are completed and spidroin proteins are cataloged, group distributions of spidroin genes were also identified in *T. antipodiana*^31^ and *Latrodectus elegans*^48^. To this end, our data fill an important information gap regarding the arrangement of spidroin genes on whole chromosomes and provide a typical entry point for further study of spider genome evolution and silk mechanical differentiation.

We found cellular clues dedicated to achieving the tri-sectional organization and functional division of the Ma gland. The Ma gland, the source of dragline silk, is composed of a folded Tail, a dilated Sac, and an elongated Duct. Our results not only define the proteomic and metabolic components of dragline silk but also trace their origins from the tri-sectional Ma gland. Previous studies have shown that there are two or three cell types that secrete spidroins in the Ma gland of orb-web spiders^14^. The cell type of the Ma gland has been debated, suggesting that it may vary between species but probably also reflecting technology difficulties in intact cell capturing after sample preparation for cellular and morphological studies. Our results obtained well-established scRNA-seq and ST evidence, indicative of ten cell types of the whole Ma gland. The fascinating single-cell spatial architecture describes the differentiation trajectory of Ma gland cells during cell state transitions, suggesting that the Sac and Duct are derived from the early cell type of Tail by cell differentiation.

We validated that the tri-sectional architecture of the silk gland was essential for silk formation. Genetic and evolutionary studies have shown that silk spun by several arthropod lineages has existed for hundreds of millions of years^49^. Numerous silk-producing animals have a variety of silk glands, which can be summed up as enlarged secretory glands connected with tiny ductal structures^35, 50^. Surprisingly, spiders and silkworms, two well-known silk producers from different classes, evolved plesiomorphic glands for silk spinning. The ASG/MSG/PSG-deficient silkworm mutant reveals three indispensable parts of the silk gland in the silk production process. Physical and material studies have shown that liquid silk dope is transformed into insoluble fiber through the combined effects of pH and ion gradients as well as extensional and shear forces as they migrate in the silk gland^3, 35^. Our work provides biological evidence for this role of the silk gland and further demonstrates a high convergence of spider Duct and silkworm ASG in several molecular functions: calcium ion binding, chitin binding, signal transduction, and *V-type H^+^-transporting ATPase* expression. Furthermore, we identified convergence silk components in spider and silkworm silks, for instance, two proteins (mucin-19 and GDH) and two major metabolites (choline and DL-malic acid). These analogous aspects implied critical processes and components essential to silk formation, which can be used as references for the optimization of Duct-based artificial spinning devices and the formula design of silk protein solutions in artificial spinning.

Based on our results, we propose a molecular model describing the comprehensive mechanism underlying dragline silk biosynthesis in the *T. clavata* Ma gland (Fig. 6): (i) The Tail, consisting of two ubiquitous cell types, is the major position for the secretion of organic acids and 13 dragline silk proteins, which consist of the inner layer of dragline silk. The presence of organic acids can affect protein solubility in water by altering conformational states^51^. In Tail, the 13 dragline silk genes are highly activated, and 47.49% of the transcripts are from these genes, among which *MaSp*-Group 1, accounting for 44.50%, is the dominant component. The relatively high chromatin accessibility (CA), along with mCG methylation, might contribute to such abundant expression. (ii) The Sac, consisting of two ubiquitous and four unique cell types, is the major position for the secretion of lipids and 28 dragline silk proteins, which consist of the middle layer of dragline silk. The lipids act as a coat wrapping around the dragline silk to regulate the water content of the silk^51^. In Sac, the 28 dragline silk genes were also highly activated, and 34.33% of the transcripts were from these genes, among which *MaSp*-Group1, *MaSp*-Group2, *MaSp9*, *SpiCE-DS1*, *SpiCE-DS2*, *SpiCE-DS4*, and *SpiCE-DS13* were dominant components. The relatively high CA, along with mCG methylation, might contribute to such abundant expression. (iii) The Duct, consisting of one ubiquitous and four unique cell types, is the major position for the secretion of chitin, cuticular protein, ions (Ca^2+^ and H^+^), and 13 dragline silk proteins. Chitin and cuticular protein construct the cuticular intima to generate shared forces^14, 52^. The ions are responsible for a multidimensional state of flux, for instance, ion exchange, pH gradient, and dehydration^11^, and the 13 dragline silk proteins consist of the outer layer of dragline silk. Unlike Tail and Sac, the transcription of dragline silk genes is less active in the Duct, and only 0.76% of the transcripts are from these genes. The relatively low chromatin accessibility (CA), along with mCG methylation, might contribute to such depressed expression. (iv) The dragline silk of *T. clavata* is composed of 10 classes of metabolites and 28 proteins. The golden pigment xanthurenic acid of dragline silk is synthesized and secreted in both the Tail, Sac, and Duct. Overall, we suspected that Tail performed a primary function of MaSps secretion. Sac, acting as a storage protein, secreted a wide range of proteins (MaSps and nonspidroin proteins), and Duct played a weak role in protein secretion but a crucial role in protein structural transition. The number of cell types in the Tail, Sac, and Duct was positively correlated with the diversity of their functions.

**Figure 6.**
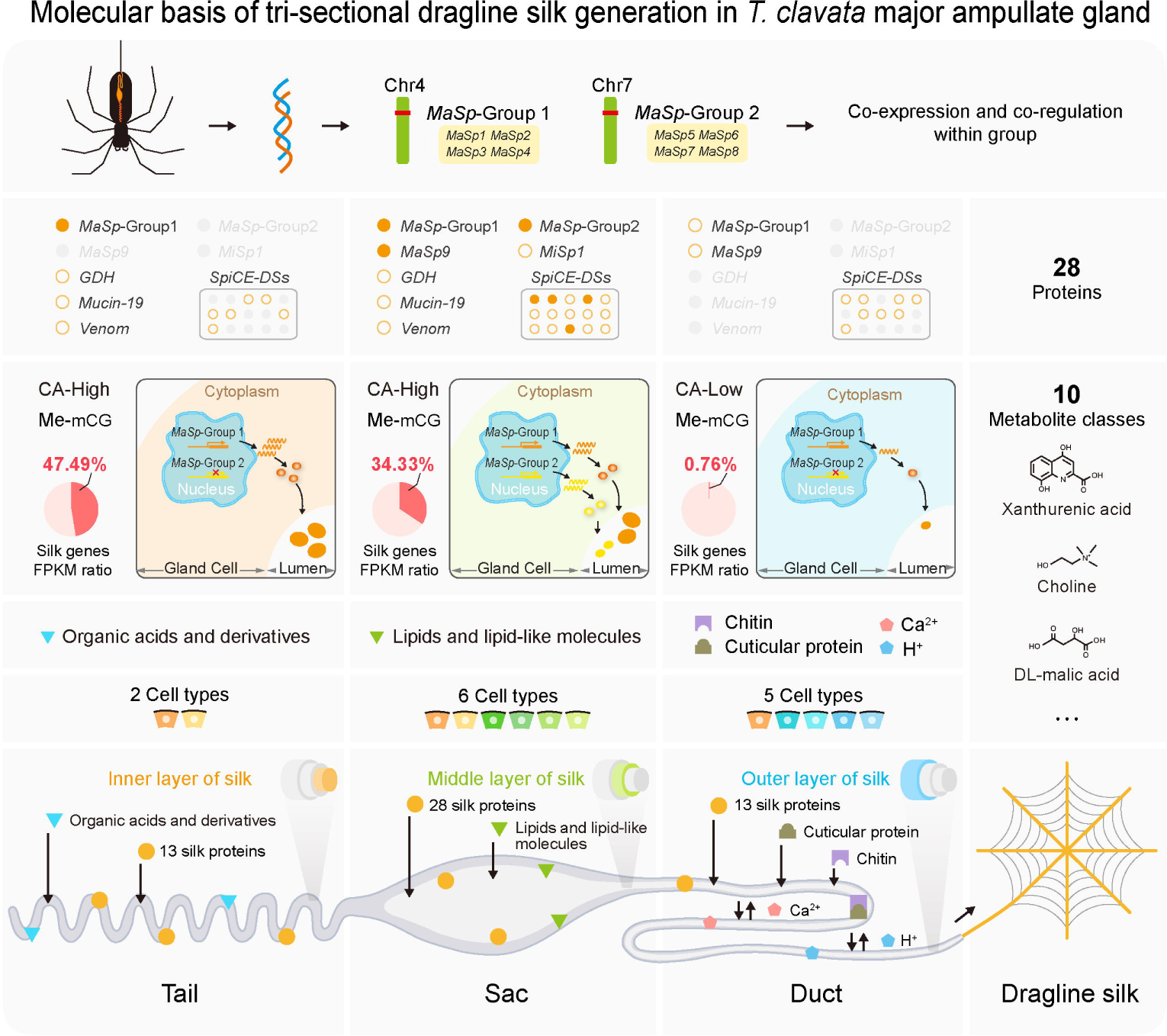
Molecular basis of tri-sectional dragline silk generation in the *T. clavata* Ma gland. Orange solid circles represent the genes with RPKM > 10,000, orange hollow circles represent the genes with RPKM < 10,000, CA: chromatin accessibility, Me: methylation.

In conclusion, our current work provides a comprehensive biological basis for dragline silk formation. We believe the chromosome-scale reference genome of the golden orb-web spider and the molecular atlas of the tri-section Ma gland presented here will facilitate an understanding of the spider silk-spinning process and further serve as a powerful platform for evolutionary studies of silk-spinning organisms. More importantly, the significant molecular characteristics and datasets in our results should ultimately pave the way for producing optimized synthetic silks through genetic and biomimetic manipulations.

## Materials and methods

### Karyotyping

Adult female *T. clavata* spiders were cultured indoors at 25 °C, and the eggs on the third day after spawning were sampled for DNA karyotyping. Approximately 20 eggs were mashed with tweezers, mixed with 1.5 mL of 0.05% colchicine for three hours, hypnotized through 0.075 mol/L KCl solution (20 min), and fixed in methanol:acetic acid (3:1) solution (30 min). The cell suspension was dropped onto precooled glass slides, flame-dried, stained with 5% Giemsa solution (50 min), rinsed with running water and air-dried. Chromosome karyotypes were observed with a 40-fold Olympus microscope.

### Genome assembly

For de novo assembly of the *T. clavata* genome, the Nanopore long reads were initially corrected using Canu v2.2^53^ with default parameters, and SMARTdenovo (https://github.com/ruanjue/smartdenovo) was subsequently employed to produce the primary contigs. Then, three rounds of contig correction were performed using Racon v1.5.0^54^. To improve the accuracy of the assembly, Pilon v1.24^55^ was utilized for contig polishing based on Illumina short reads. The paired Hi-C short reads were primarily trimmed by Hic-pro v2.10^56^, and the optimized reads were mapped to the draft contig with Juicer v1.6.2^57^ using 3D-DNA v180419^58^ for pseudochromosome construction and manual corrections with the Juicebox v1.9^57^ tool.

### Genome annotation

#### Repeat annotation

A custom *T. clavata* repeat database was de novo identified by RepeatModeler v2^59^ and LTR_FINDER v1.06^60^. Then, the repeat sequences generated above were imported into RepeatMasker v4.05^61^ to search for transposable elements (TEs) in the genome. Tandem Repeats Finder v4.09^62^ was used to find tandem repeats.

#### Gene annotation

Gene structure prediction was based on homology evidence and de novo prediction. The evidence files contain RNA-seq data (17 tissues RNA-seq), Iso-seq data, and protein sequences of close species (Four species: *A. ventricosus*, *A. bruennichi*, *T. antipodiana*, and *T. clavipes*). AUGUSTUS^63^ (http://bioinf.uni-greifswald.de/) software was used for gene de novo prediction. Evidence-based gene annotations were produced in the MAKER2^32^ pipeline with default parameters. Then, we integrated the MAKER and de novo results based on the following screening criteria: the proportion of repeat sequences was less than 50%, the protein sequence length was greater than 50 aa, the FPKM value was greater than 0.1 in at least ten samples, and the sequence was supported by Iso-seq evidence (identity > 95%) or at least one closely related species (E-value > 1e^-5^, identity > 50%).

### Spidroin analysis

#### Spidroin identification

A total of 443 redundant spidroin sequences (Supplementary Table 6) were downloaded from the NCBI protein database (https://www.ncbi.nlm.nih.gov/protein), using all protein sequences of *T. clavata* to align these spidroins by blastp (E-value < 1e^-10^), and 128 nonrepetitive hits were primarily identified. To further identify reliable spidroins, we double-checked that each sequence must include typical repeats and N- or C-termini, and the final 28 putative spidroins were recognized.

#### Artificial full-length spidroin assembly

Briefly, 1) 79 complete spidroin sequences, 48 N-terminal, and 22 C-terminal sequences (Supplementary Table 7) were mapped to the *T. clavata* genome to generate the GFF track files using the genBlast v138^64^ tool; 2) the bam files of silk gland Iso-seq and the bam files of silk gland RNA-seq data were extracted as expression evidence tracks; 3) the GFF3 files of 28 spidroins were extracted as one track. 4) The track files of the previous three steps were imported into IGV v2.9.4^65^, and the integrity of each spidroin was manually checked. 5) The incomplete spidroin sequences were repaired by cutting the corresponding DNA sequence in the genome based on frameshift translation and prediction using the online tool AUGUSTUS^63^ (http://Augustus.gobics.de/).

#### Spidroin motif analysis

Amino acid content and motifs were counted using a Perl script. The motifs included (GA)n, (A)n, GGX, XQQ, and GPGXX. O-glycosylation was predicted with the NetOGlyc v4.0^66^ tool.

### Proteomics

The dragline silk of spider (*T. clavata*) and cocoon silk of silkworm (*B. mori*) were cut with scissors. First, silk (20 mg per sample) was dissolved in 500 μL of 9 M LiSCN, and the supernatant was collected after centrifugation (10,000 × g, 10 min). The protein concentration was measured by the Bradford protein assay (Beyotime, China). Then, the samples were diluted 10-fold in 8 M urea solution, mixed with 5 × SDS–PAGE buffer, and separated using a NuPAGE 4–12% Bis-Tris protein gel (Thermo Fisher Scientific, USA). Samples were digested using trypsin, and peptides were measured by LC–MS/MS on a Q Exactive HF-X (Thermo Fisher). The identification and iBAQ of peptides were analyzed with MaxQuant v1.3.0.5^67^.

### Metabolomics

Shredded dragline silk and cocoon silk samples were used for metabolite extraction. The silk sample was placed in a tube, homogenized in 500 μL 80% methanol, kept on ice for 10 min, and centrifuged at 1,5000 × g for 20 min at 4 °C. The supernatant (250 µL per sample) was collected for LC–MS/MS experiments on a HypesilGold column (C18). The metabolite identities and quantities were confirmed by compound Discoverer (CD) software analysis, which also performed alignment at the mzCloud (https://www.mzcloud.org/), mzVault, and MassList databases. Furthermore, the KEGG^68^ (https://www.kegg.jp/), HMDB^69^ (https://hmdb.ca/metabolites), and LIPIDMaps^70^ (http://www.lipidmaps.org/) databases were utilized for metabolite annotation.

### Methylation analysis

#### Whole-genome bisulfite sequencing (WGBS)

The Tail, Sac, and Duct of the Ma gland were used for WGBS. First, genomic DNA was extracted and fragmented to a mean size of 250 bp by sonication using a Bioruptor (Diagenode, Belgium), followed by adding an “A” base to the 3’ terminal ends of DNA fragments and ligating methylated adaptors to both ends of DNA fragments. The EZ DNA Methylation-Gold Kit (ZYMO) was used for bisulfite conversion of ligated DNA. Products were purified using a QIAquick Gel Extraction kit (Qiagen) and amplified by PCR.

Finally, libraries were sequenced on the Illumina HiSeq 2500 platform.

#### Methylation level calculation

The clean reads were mapped to the reference genome by using BSMAP (-v 8 -z 33 -p 4 -n 0 -w 20 -s 16 -f 10 -L 100) software after removing the adaptor and low-quality reads (N > 10% and Q < 20). Methylation levels were calculated through the beta value: beta = M/(M + U), where M and U represent methylated and unmethylated signal values. The bed file of methylation was converted to bigwig format by the “bedGraphToBigWig” command line and visualized in IGV^65^.

### Assay for transposase-accessible chromatin

#### ATAC-seq library preparation and sequencing

The Tail, Sac, and Duct of the Ma gland were used for ATAC sequencing. Samples were ground individually in liquid nitrogen and transferred to precooled cell lysate at 4 °C for 10 min. The cell pellets were retained when the supernatant was removed after being centrifuged (500 × g, 4 °C, and 10 min), washed with precooled buffer solution, and centrifuged (500 × g, 4 °C, and 10 min) to collect cell nuclei. The cell nucleus was mixed with Tn5-transposase and kept for 30 min at 37 °C, followed by DNA fragment collection, PCR amplification, and sequencing on the Illumina PE150 platform.

#### Chromatin accessibility detection

The clean reads were mapped to the genome using Bowtie2^71^ software, converted from.sam format to.bam format, and filtered reads by using Samtools v0.1.19^72^ with parameters (view -b -f 2 -q 30). The peaks of chromatin accessible regions were identified from each sample separately using MACS2^73^ (--shift -100 --extsize 200 --nomodel -B -q 0.05). The “intersect” command of BEDTools v2.26.0^74^ was used to identify shared peaks between replicate samples. Heatmaps were displayed using the “plotHeatmap” function of deepTools v3.5.1^75^. The significantly enriched motifs were discovered by the HOMER^76^ tool.

### Whole-transcriptome analysis

Total RNA was extracted from the Tail, Sac, and Duct of the Ma gland with TRIzol reagent, followed by library construction for lncRNAs and *miRNAs*.

#### LncRNA sequencing and analysis

Total RNA samples were incubated with DNase I to digest DNA fragments and RNase H to remove rRNA. After purification, the cDNA was synthesized with the base “A” added to the 3’ end and ligated to the adapter, followed by UDG digestion of the second strand and PCR amplification. The constructed library was sequenced on DNBSEQ. The clean reads were mapped to the reference genome using HISAT2^77^ (--phred64 --sensitive --no-discordant --no-mixed -I 1 -X 1000), and the transcripts were assembled using StringTie^78^ (-f 0.3 -j 3 -c 5 -g 100 -s 10000 -p 8) and compared with known mRNAs using the cuffcompare (-p 12) program of Cufflinks^79^. Then, cooperative data classification (CPC), txCdsPredict, and coding noncoding index (CNCI) methods were used to distinguish the mRNAs and lncRNAs. Cis and trans methods were used to evaluate the target mRNAs of lncRNAs. Cis refers to the lncRNAs located 10 kb upstream or 20 kb downstream of the target mRNA. The binding energy (BE) of lncRNAs and mRNAs was analyzed by RNAplex^80^, and if BE < -30, it was considered trans.

#### Small RNA sequencing and analysis

Total RNA was purified and isolated using PAGE electrophoresis and excised to obtain 18-30 nt RNA. The 3’ end of the small RNA fragment was ligated to the 5-adenylated and 3-blocked adaptor, and the unique molecular identifiers (UMI)-labeled primer was added to hybridize with the 3’ adaptor. Then, the 5’ adaptor was hybridized with the 5’ end of the UMI label. The cDNA strand was synthesized with UMI-labeled primers, and highly sensitive polymerase was used to amplify the product. We used Bowtie2^71^ (-q -L 16 --phred64 -p 6) to align the clean reads to the reference genome and the small RNA database Rfam^81^ (using cmsearch with the parameters --cpu 6 --noali). To identify miRNAs, which were annotated and predicted via the miRBase^82^ database (https://www.mirbase.org/) and miRDeep2^83^ tool, their targets were predicted using miRanda^84^ (-en -20 -strict) and RNAhybrid^85^ (-b 100 -c -f 2,8 -m 100000 -v 3 -u 3 -e -20 -p 1) between miRNAs and lncRNAs/mRNAs.

### Single-cell transcriptome

#### Sample processing and sequencing

Freshly dissected Ma gland was washed using culture medium, cut into small pieces and digested in a constant temperature incubator for 45 min. After digestion, the medium was filtered through a 40 μm cell sieve and washed more than two times (centrifuge for 5 min at 300 × g at 4 °C). Finally, the activity of the cell suspension greater than 85% and the agglomeration rate less than 5% were used for library construction and sequencing. The library was constructed according to the manufacturer’s instructions (10 × Genomics) on a 10 × Chromium Single Cell 3’ Platform.

#### Data analysis

The raw data were imported into the cellranger v4.0.0 pipeline to obtain the expression matrices per cell and per gene. To remove low-quality cells, cell numbers and gene UMI counts within the range of the median ± 2-fold median absolute deviation (MAD) were retained for further analysis. Additionally, doublets (≥ 2 cells in one oil droplet) were removed using DoubletFinder v2.0.3^86^. A total of 9,349 high-quality cells were used to analyze clustering and expression based on the Seurat v4.0.6^87^ package. The developmental trajectories of all cell clusters were predicted by the Monocle v2.22.0^88^ package.

### Spatial Transcriptomics

#### Sample processing and sequencing

Fresh Ma gland was frozen with isopentane in liquid nitrogen and stored in an airtight container at -80 °C after being embedded with OCT. Embedded frozen tissue was sectioned, and sections of acceptable RNA quality (RIN > 7.0) were used for further experiments. Then, the sections were placed on Visium Spatial slides, fixed with methanol, and stained with hematoxylin and eosin. The polyadenylated mRNA was captured by the primers of spots. The RT Master Mix reverse transcription reagent was added to permeate tissue sections, and second strand synthesis was performed on the slide by adding the Second Strand Mix. The cDNA was denatured from each capture zone and transferred to the corresponding tube for amplification, library construction, and sequencing on the Illumina NovaSeq600 platform.

#### Data analysis

After removing adapters and low-quality (N > 3, Q < 5, and base ratio ≥ 20%) reads, the spaceranger v1.3.1 mkref and count programs were used for data preprocessing. Then, spot clustering was analyzed using the Seurat v4.0.6^87^ package.

### Convergent evolution analysis

To compare whether there was a convergent evolution between the silk gland of silkworm and the Ma gland of spider. We compare them in five aspects (see supplementary note for details): 1) the morphological structure of silk glands; 2) the scanning electron microscope (SEM) microstructure of silk, dragline silk, and cocoon silk; 3) the expression convergence of silk gland genes; 4) protein components in dragline silk and cocoon silk; and 5) metabolite components in dragline silk and cocoon silk. The orthologous genes between silkworm and spider were identified using the blastp^89^ (E-value < 1e^-5^) program.

### Data availability

High-throughput sequencing raw data in this project were deposited into the CNGB Nucleotide Sequence Archive (CNSA) of the China National GeneBank DataBase (CNGBdb, http://db.cngb.org) and are available through BioProject ID CNP0002864. The genome assembly and gene annotation of *Trichonephila clavata* are also available on SpiderDB (https://spider.bioinfotoolkits.net/).

## Author contributions

W.H., A.J., Q.X., and Y.W. conceptualized the project. Y.W. supervised the project. W.H., A.J., Z.Z., J.S., T.Y., T.X., Q.L., T.Y., J.Z. and Y.W. collected samples for multiomics sequencing. A.J. designed and implemented the genome assembly pipeline and performed data analysis. W.H. and A.J. drafted the manuscript. F.L., Y.L. and Z.J. generated the data and built the database. W.H. and S.M. conducted the experiments. Y.W. finalized the manuscript with input from all authors. All authors read and approved the final manuscript.

## Acknowledgments

We thank Daojun Cheng at Southwest University and Yi Zou at Southwest University for critical reading of the manuscript. We thank Yazhou Li at Sichuan Agricultural University for providing help in karyotyping. We are very grateful to many anonymous reviewers for testing the database and offering valuable comments.

## Supplementary material

**Supplementary Figure 1.**
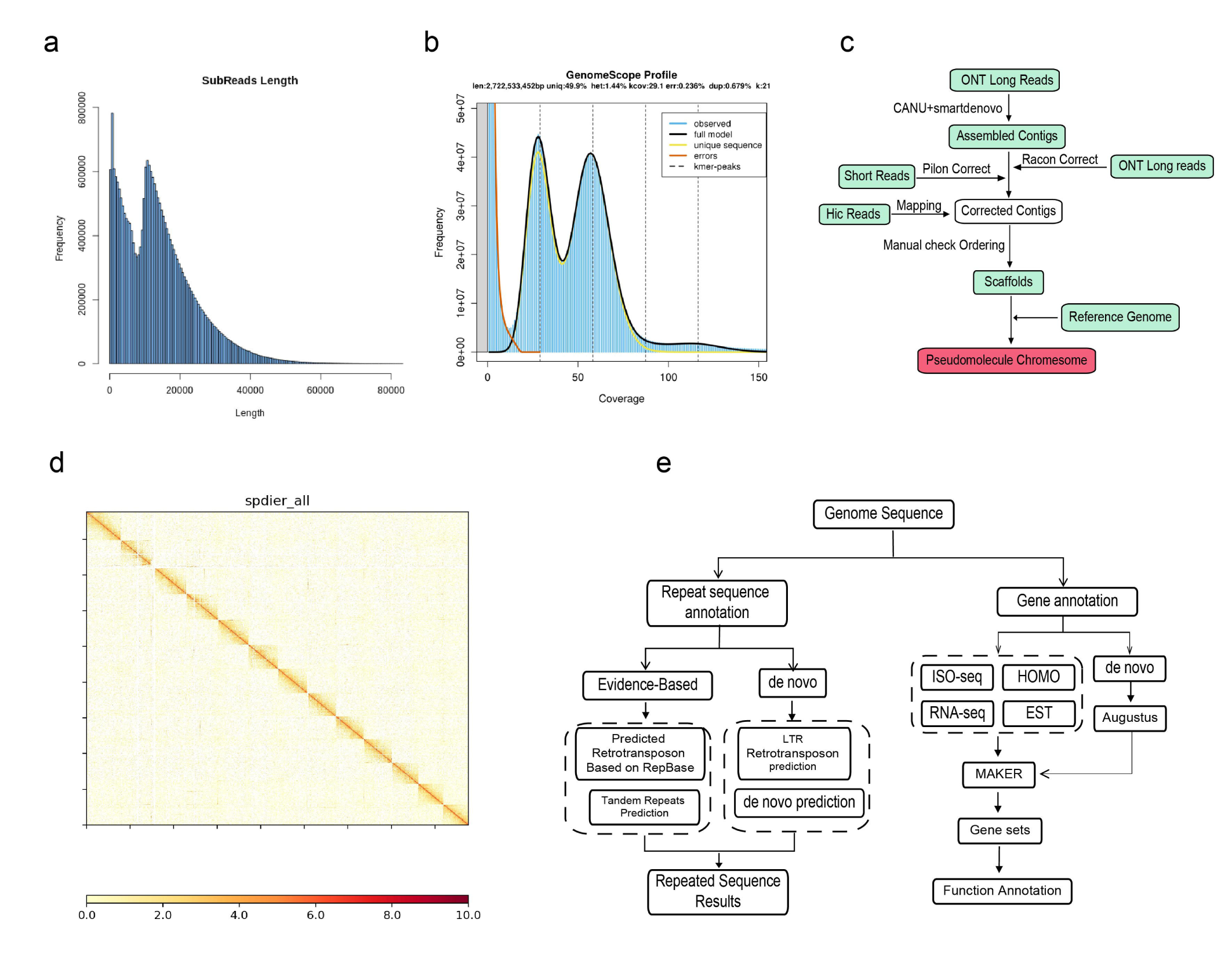
Brief information of genome assembly and annotation for female *T. clavata*. **a**, The peaks of the genome survey for k = 21. **b**, Histogram of ONT read distribution. **c**, Brief pipeline of genome assembly. **d**, Hi-C heatmap of pseudochromosome construction. **e**, Brief pipeline of genome annotation.

**Supplementary Figure 2.**
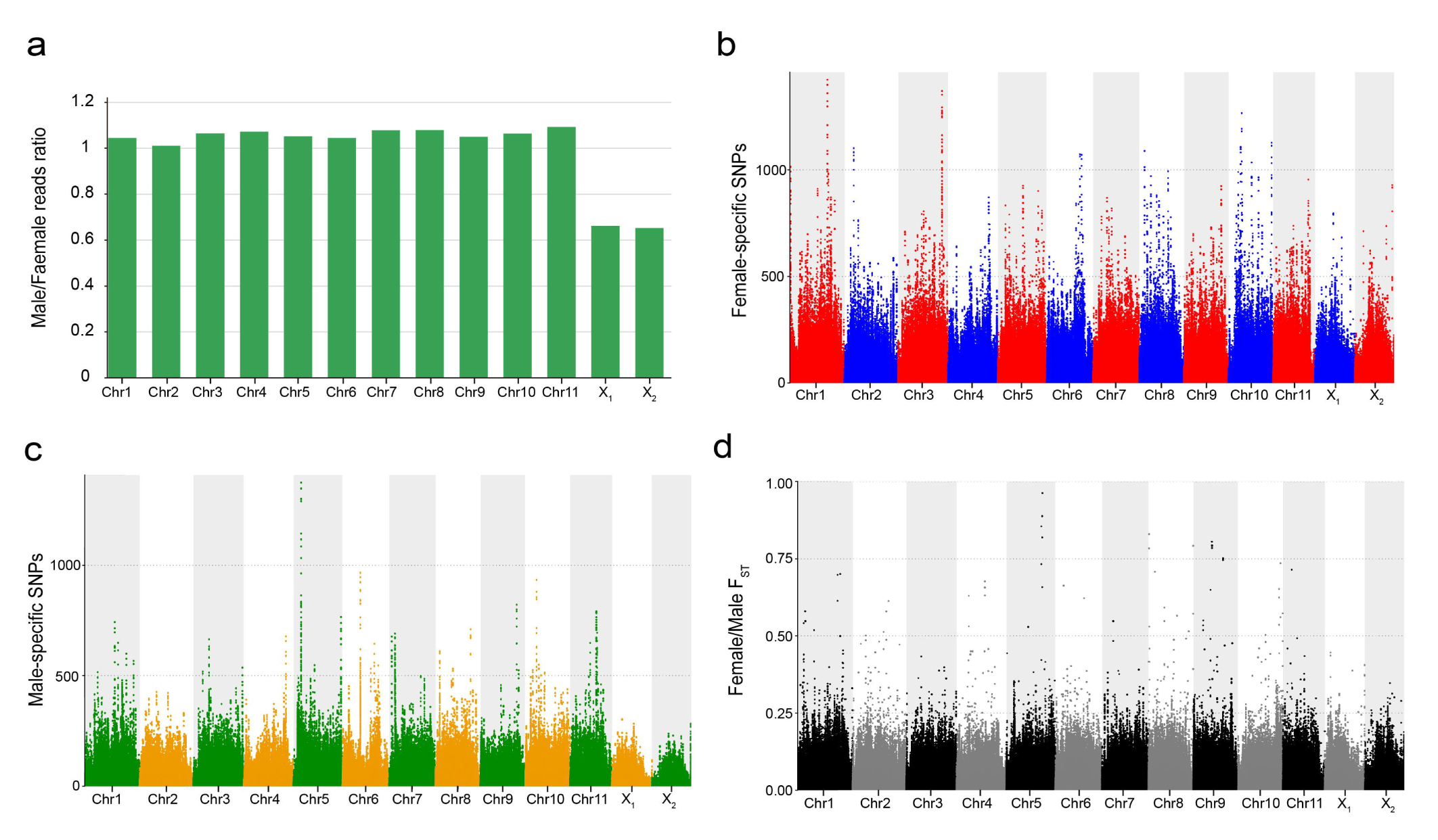
Sex chromosome identification and analysis for *T. clavata*. **a**, The mapped read ratio of pool-seq in each chromosome. Manhattan plots for **b**, Female-specific SNPs, **c**, Male-specific SNPs, and **d**, Female/Male F_ST_.

**Supplementary Figure 3.**
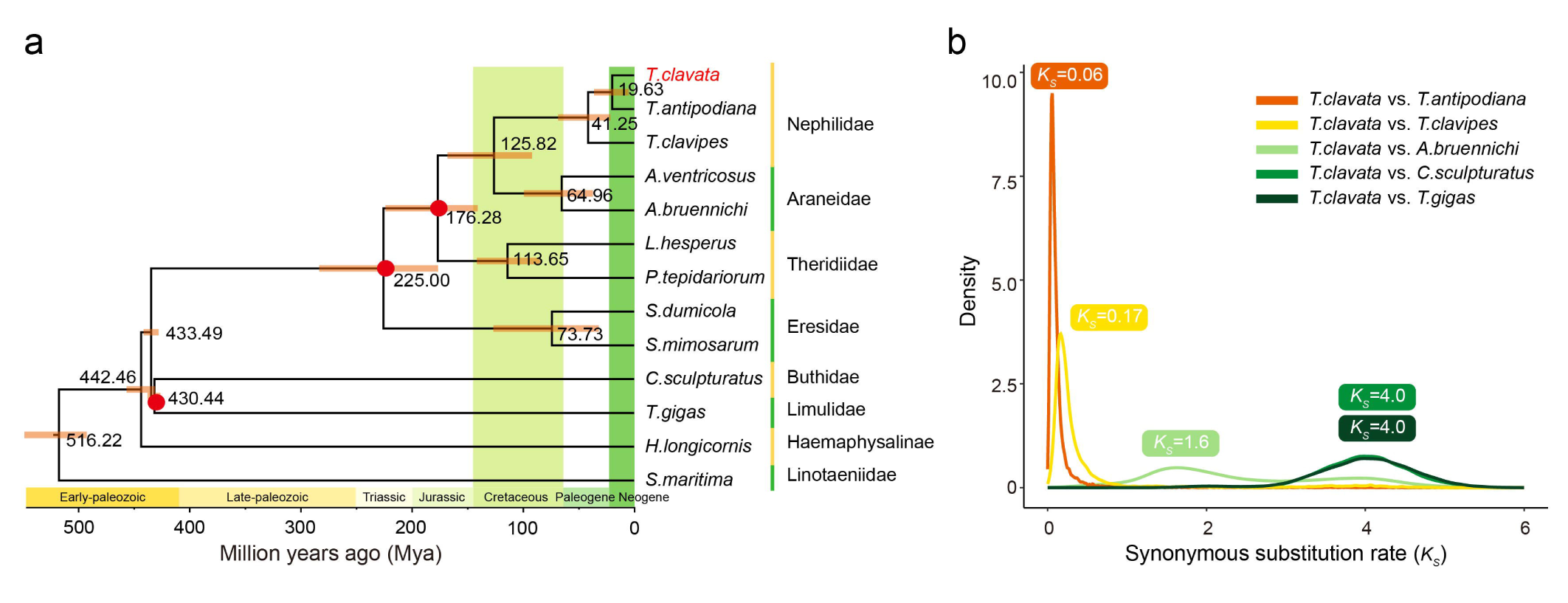
Evolutionary features in the genome. **a**, Phylogenetic tree for 13 species. The red circles represent the time correction points from the timetree (http://www.timetree.org/), and the numbers on the nodes represent the divergence time. **b**, The distribution of the synonymous substitution rate (*K_s_*) of homologous genes between *T. clavata* and five arthropod species (*T. antipodiana*, *T. clavipes*, *A. bruennichi*, *C. sculpturatus,* and *T. gigas*). Five peaks (*K_s_* = 0.06, 0.17, 1.6, 4.0, and 4.0) of *K_s_* distribution indicate the divergences of *T. clavata* and *T. antipodiana*, *T. clavipes*, *A. bruennichi*, *C. sculpturatus,* and *T. gigas*.

**Supplementary Figure 4.**
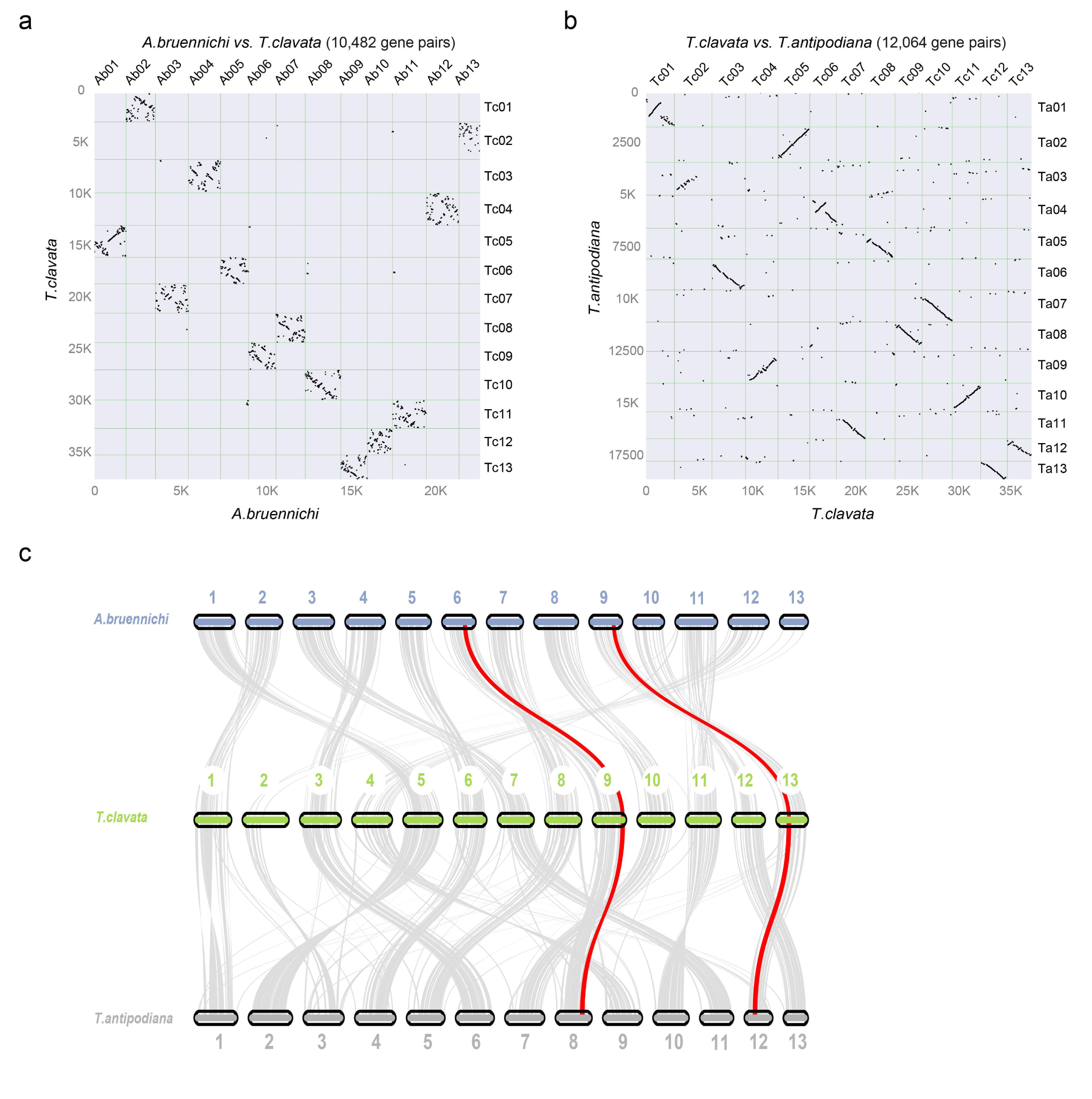
Collinearity relationships of genes. **a**-**b**, The dot plot of orthologous gene pairs for *A. bruennichi* and *T. clavata, T. antipodiana* and *T. clavata*, respectively. **c**, Gene collinearity between *A. bruennichi*, *T. antipodiana,* and *T. clavata*; the red lines represent *Hox* clusters.

**Supplementary Figure 5.**
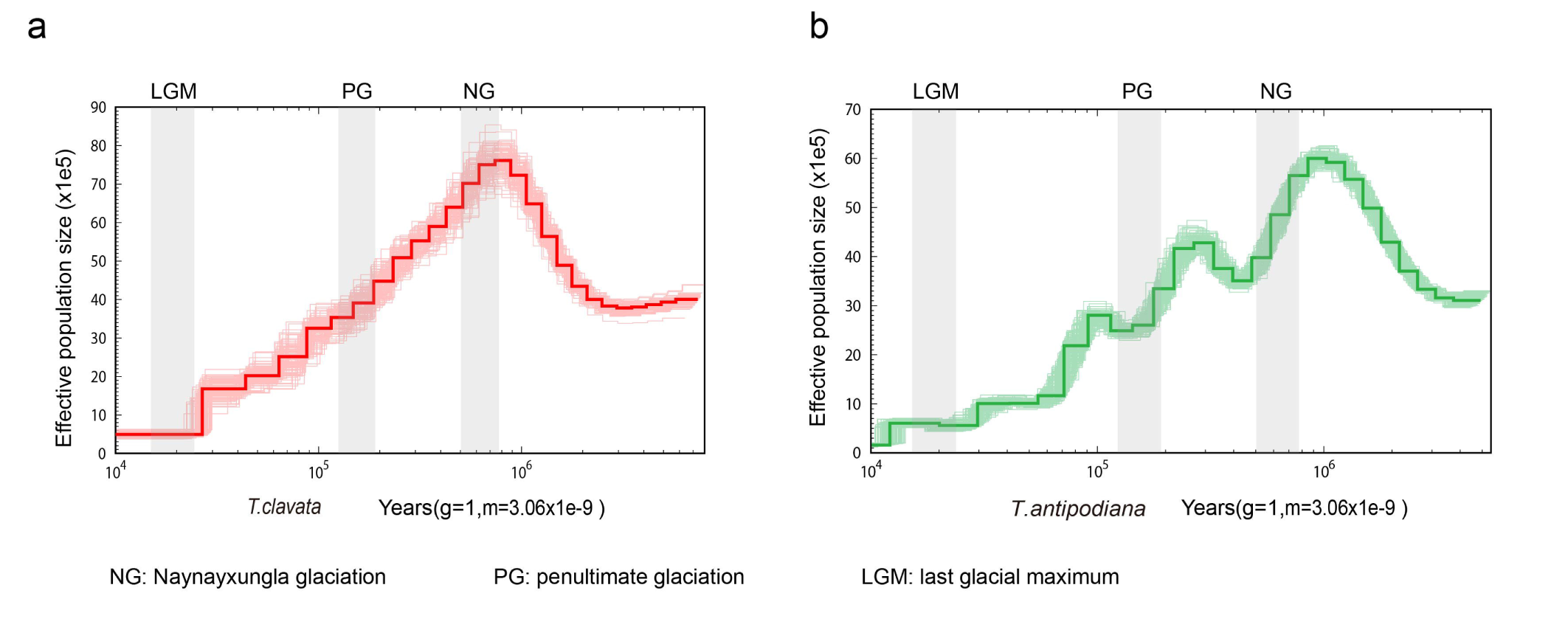
Demographic history. **a**-**b**, Population history of *T. clavata* and *T. antipodiana*. Effective population size analysis revealed that the *T. clavata* and *T. antipodiana* spiders experienced population expansion at approximately 1,000,000 years.

**Supplementary Figure 6.**
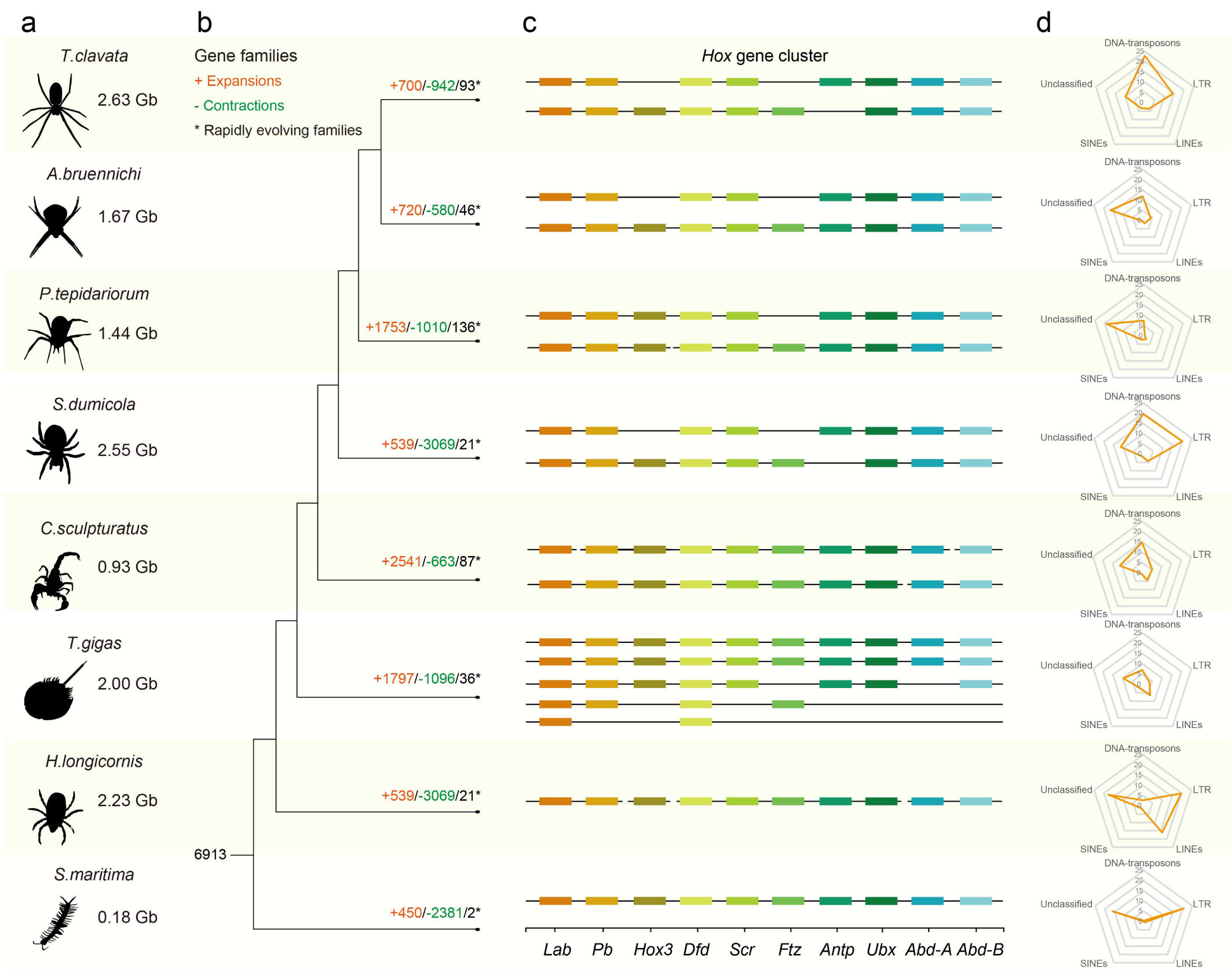
**a**, Mode charts and genome size statistics for eight species. **b**, Expansion, contraction, and rapidly evolving gene families. **c**, *Hox* cluster distribution. **d**, Radar chart of TE content.

**Supplementary Figure 7.**
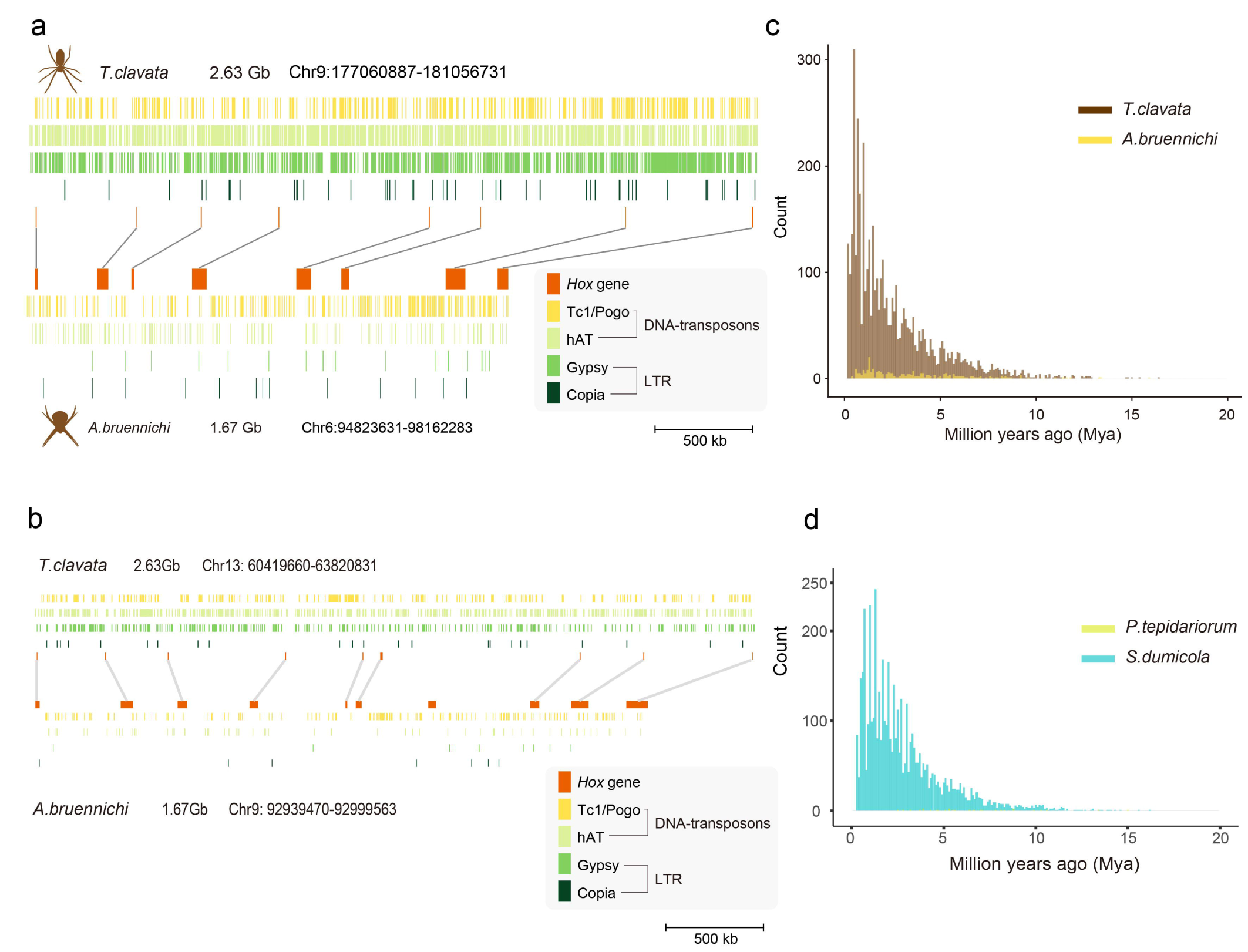
Transposon distribution. **a**-**b**, The insert distribution of the DNA (Tc1/Pogo and hAT) and RNA transposons (LTR: Gypsy and Copia) in the *Hox* cluster of *T. clavata* and *A. bruennichi*. **c**-**d**, Histogram of LTR insert time for *T. clavata* and *A. bruennichi*, *P. tepidariorum* and *S. dumicola*.

**Supplementary Figure 8.**
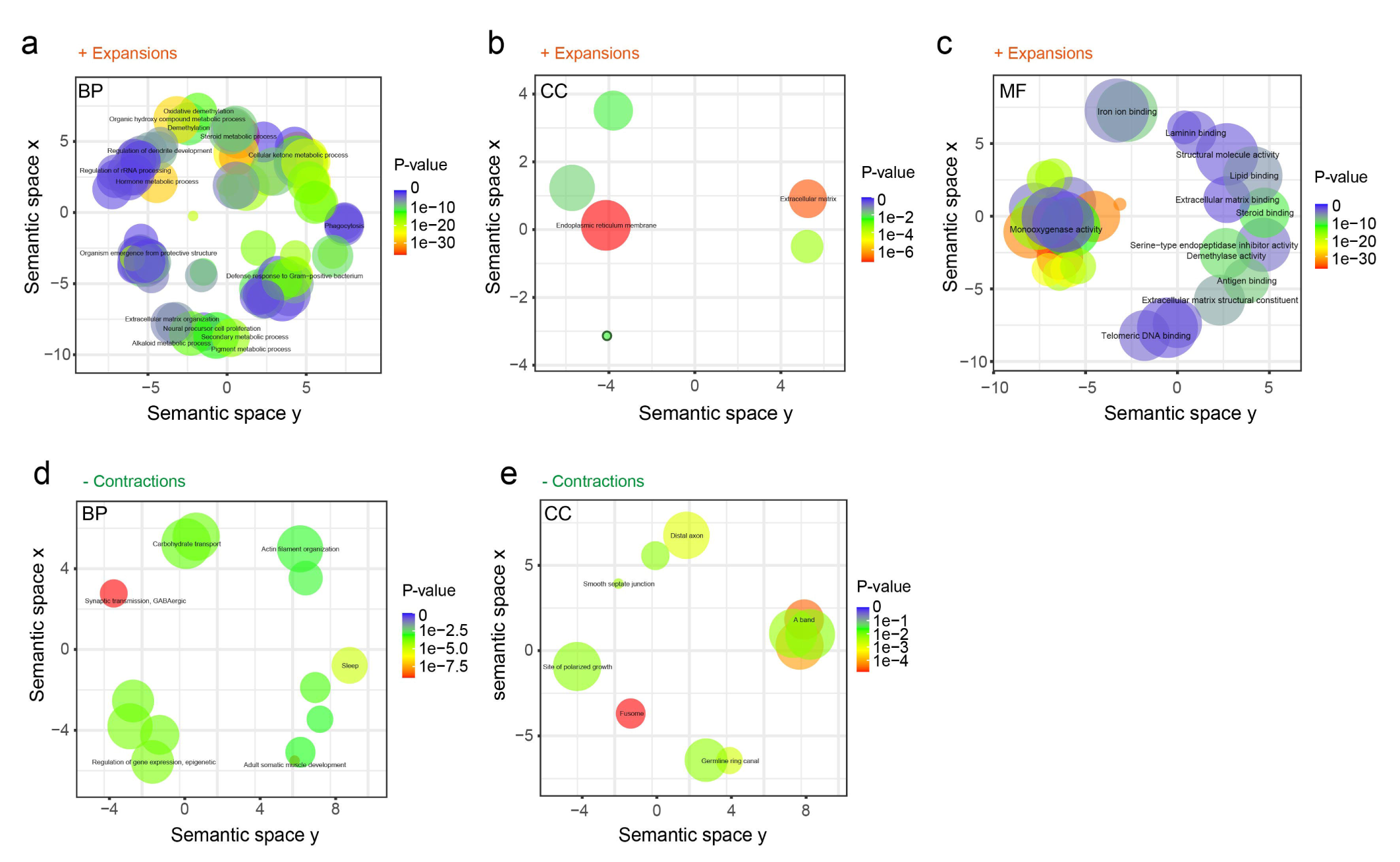
GO annotations for expansion and contraction gene families in *T. clavata*. **a-c**, Expanded gene families were enriched in the GO terms of biological process (BP), cellular component (CC), and molecular function (MF). **d-e**, Contracted gene families were enriched in the GO terms of BP and CC but not enriched in MF.

**Supplementary Figure 9.**
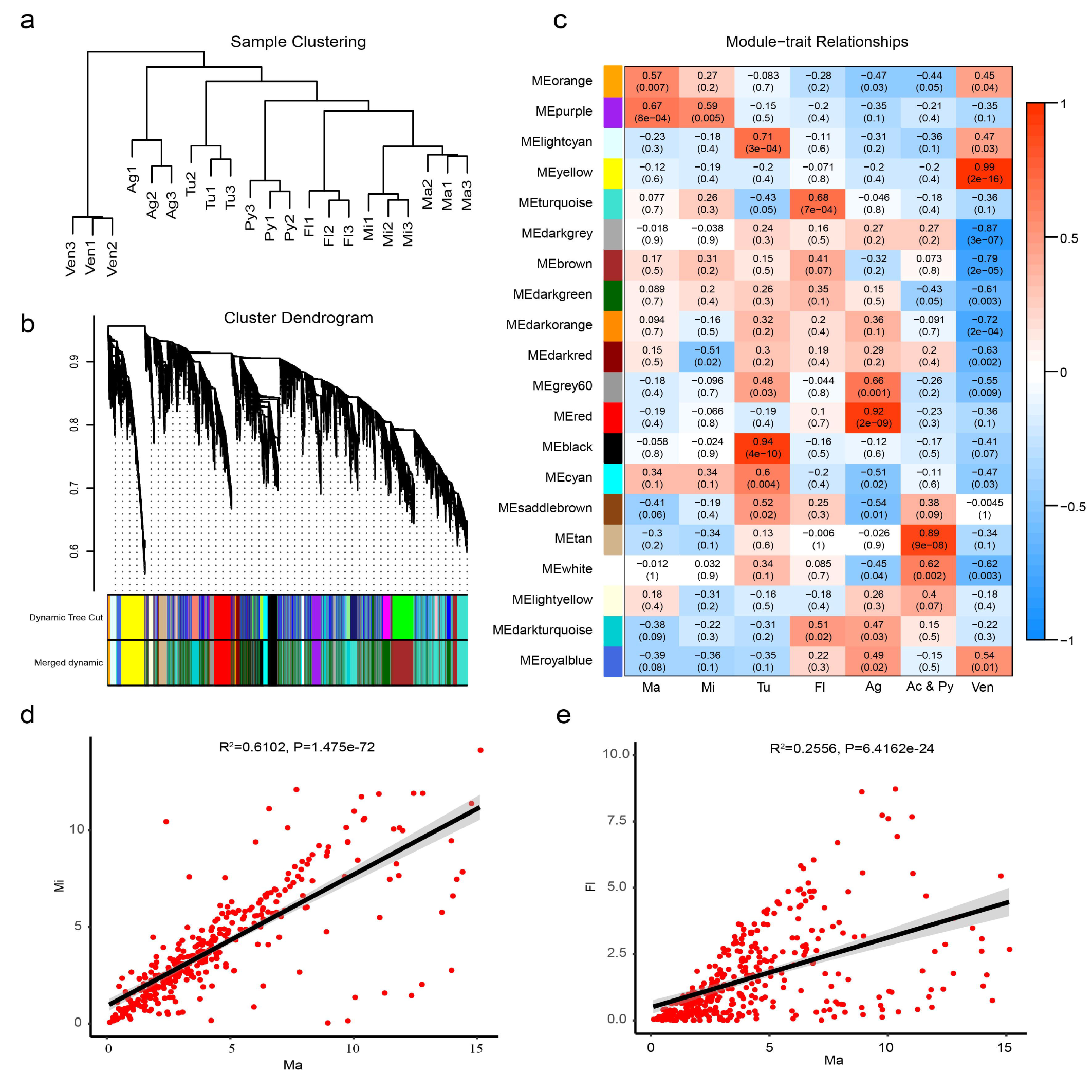
WGCNA for six silk glands and one venom gland. **a**, Sample clustering. **b**, Cluster dendrogram of genes. **c**, Module-trait relationships. **d-e**, The correlation relationships of the genes of the purple module between Ma and Mi or Fl. The results indicated that Ma was closer to Mi than to other silk gland tissues.

**Supplementary Figure 10.**
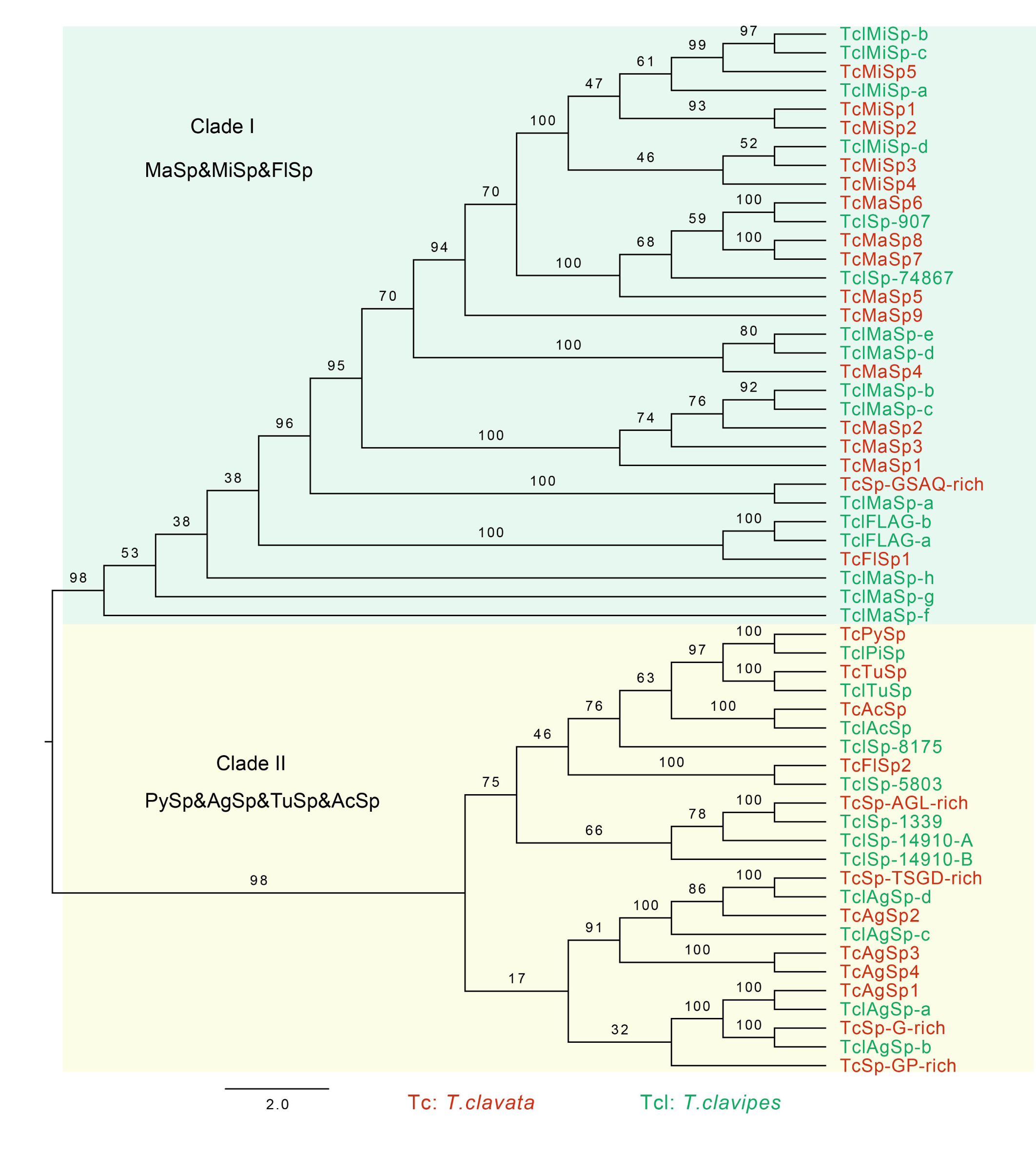
Phylogenetic relationships of 54 spidroins. This tree was constructed using protein sequences. FlSp and FLAG represent flagelliform gland spidroins, and PiSp and PySp represent pyriform gland spidroins.

**Supplementary Figure 11.**
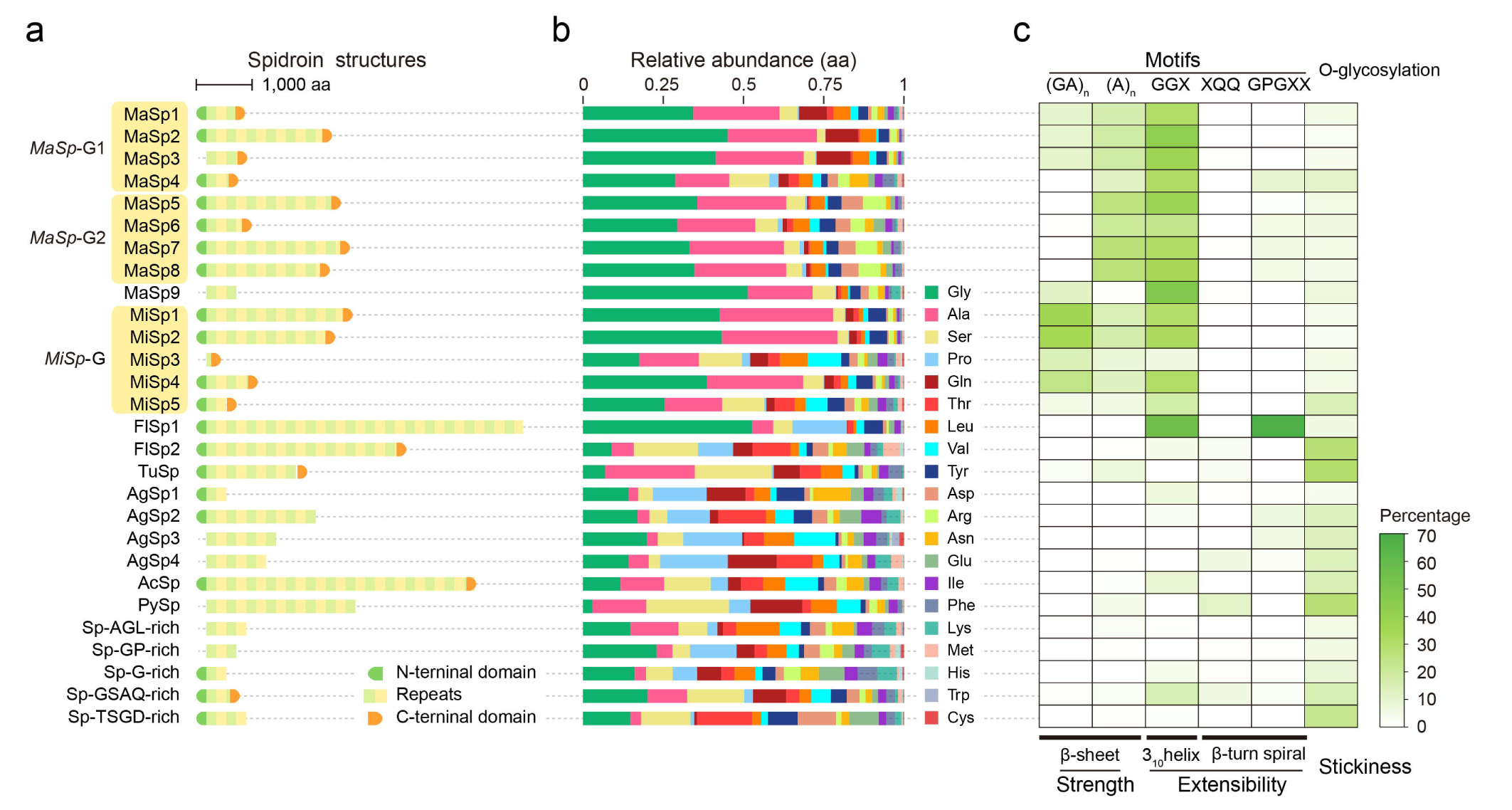
Spidroin sequence characteristics. **a**, Spidroin structure columns showing the N/C-terminal (green and orange box) and repeat domains. Each structure is drawn to scale. **b**, Amino acid content of spidroins. **c**, Heatmap showing the variety of repetitive motifs in spidroins.

**Supplementary Figure 12.**
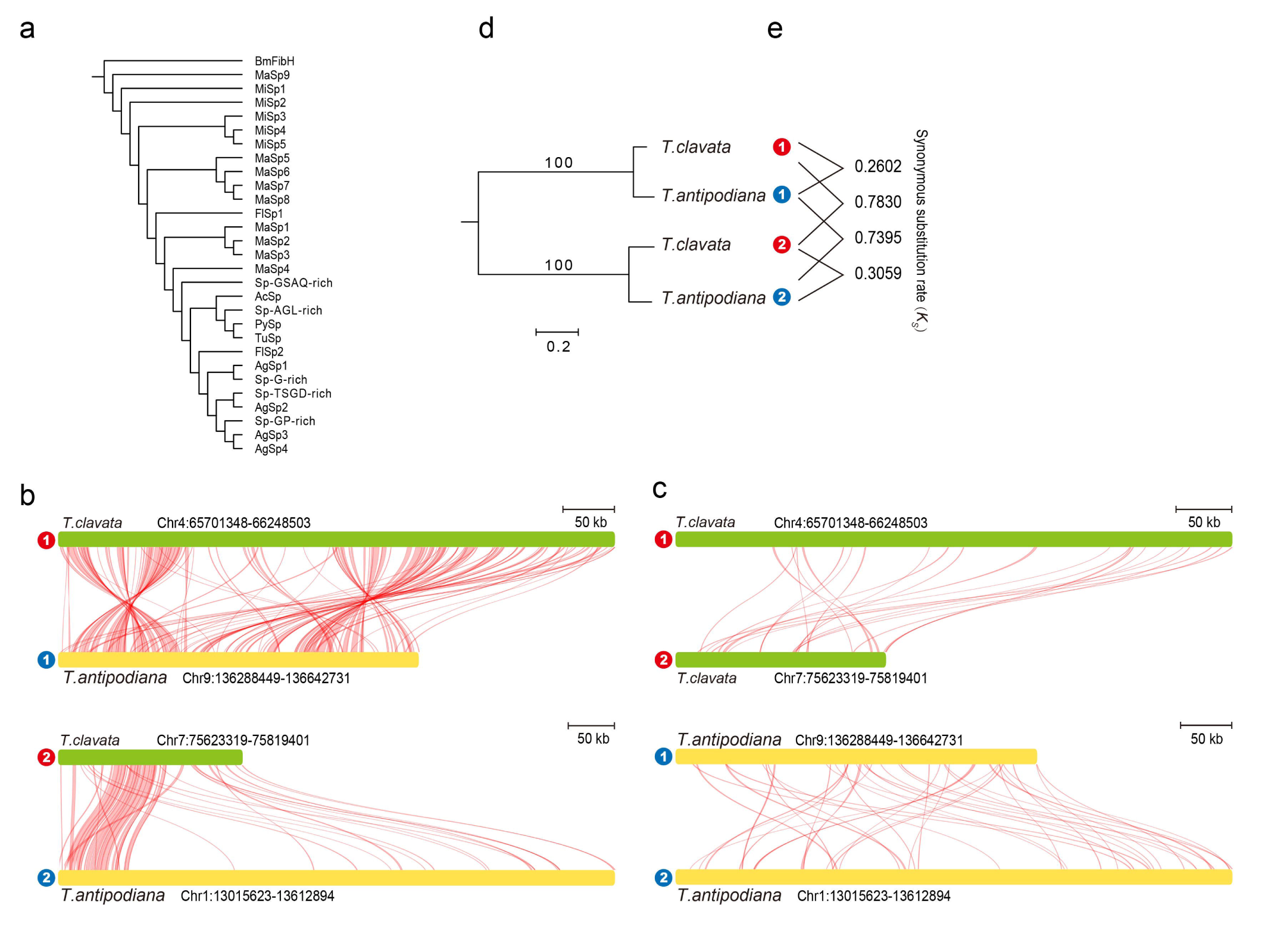
Spidroin evolution. **a**, Phylogenetic relationships of 28 spidroins and BmfibH (*B. mori*) as an outgroup. **b**-**c**, Collinearity for *MaSp*-Group 1 and -Group 2. The results showed that collinearity was higher between species than within species. **d**-**e**, The evolutionary relationships and *K_s_*values of *MaSp* clusters for *T. antipodiana* and *T. clavata*. The phylogenetic relationship within species was far greater than that between species for *MaSp*-Group 1 or MaSp-Group 2 and indicated that *MaSp*-Group 1 has a common ancestor. *K_s_*values also showed similar results (*Tc*-*MaSp*-Group 1 vs. *Ta*-*MaSp*-Group 1: 0.2602 was less than *Tc*-*MaSp*-Group 1 vs. *Tc*-*MaSp*-Group 2: 0.7830, also less than *Ta*-*MaSp*-Group 1 vs. *Ta*-*MaSp*-Group 2: 0.7395).

**Supplementary Figure 13.**
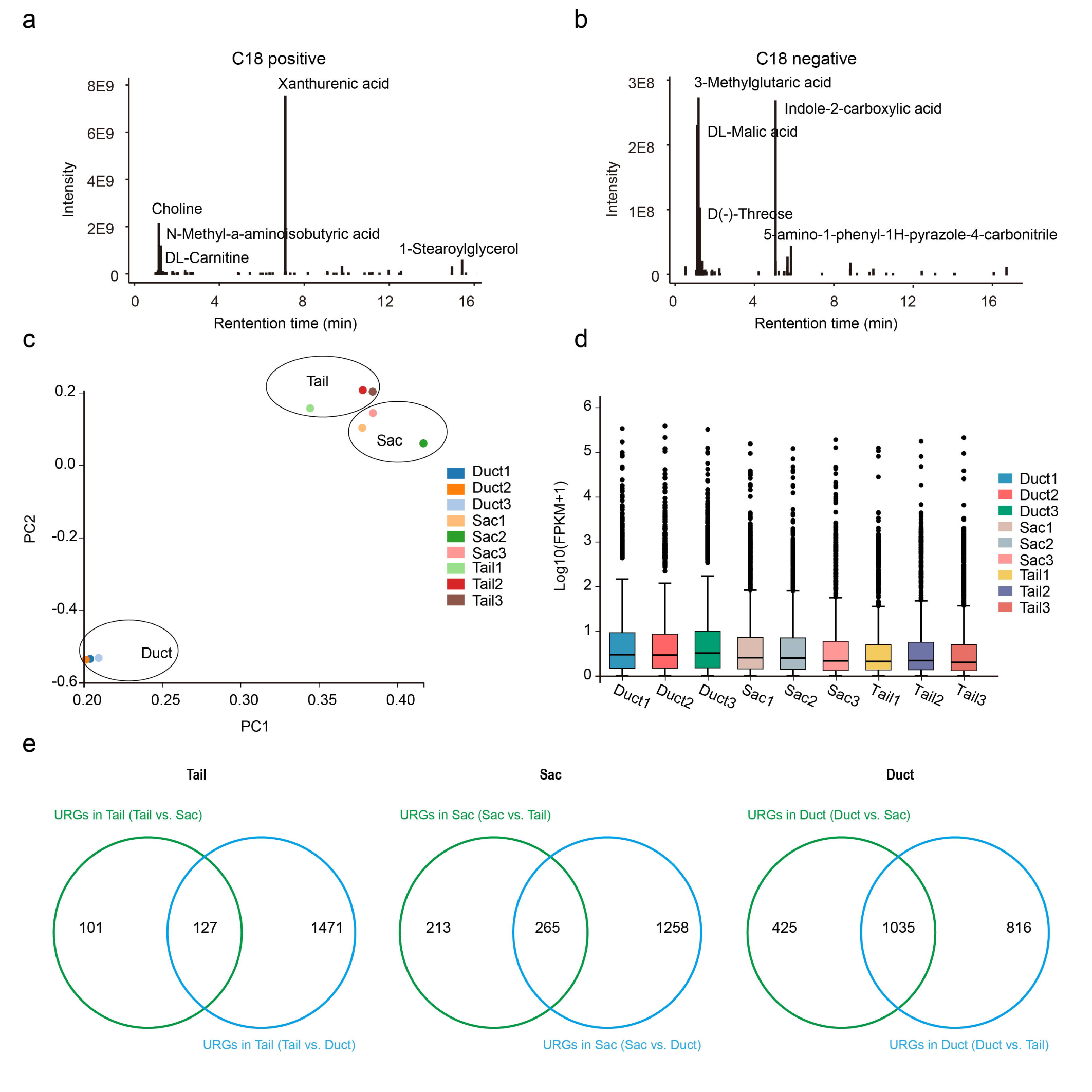
**a-b**, LC–MS analyses of the metabolites in dragline silk. **c-d**, Principal component analysis plot and boxplot of FPKM expression levels of nine samples. **e**, Flowchart of the screening of unique genes in the Tail, Sac and Duct of the MA gland.

**Supplementary Figure 14.**
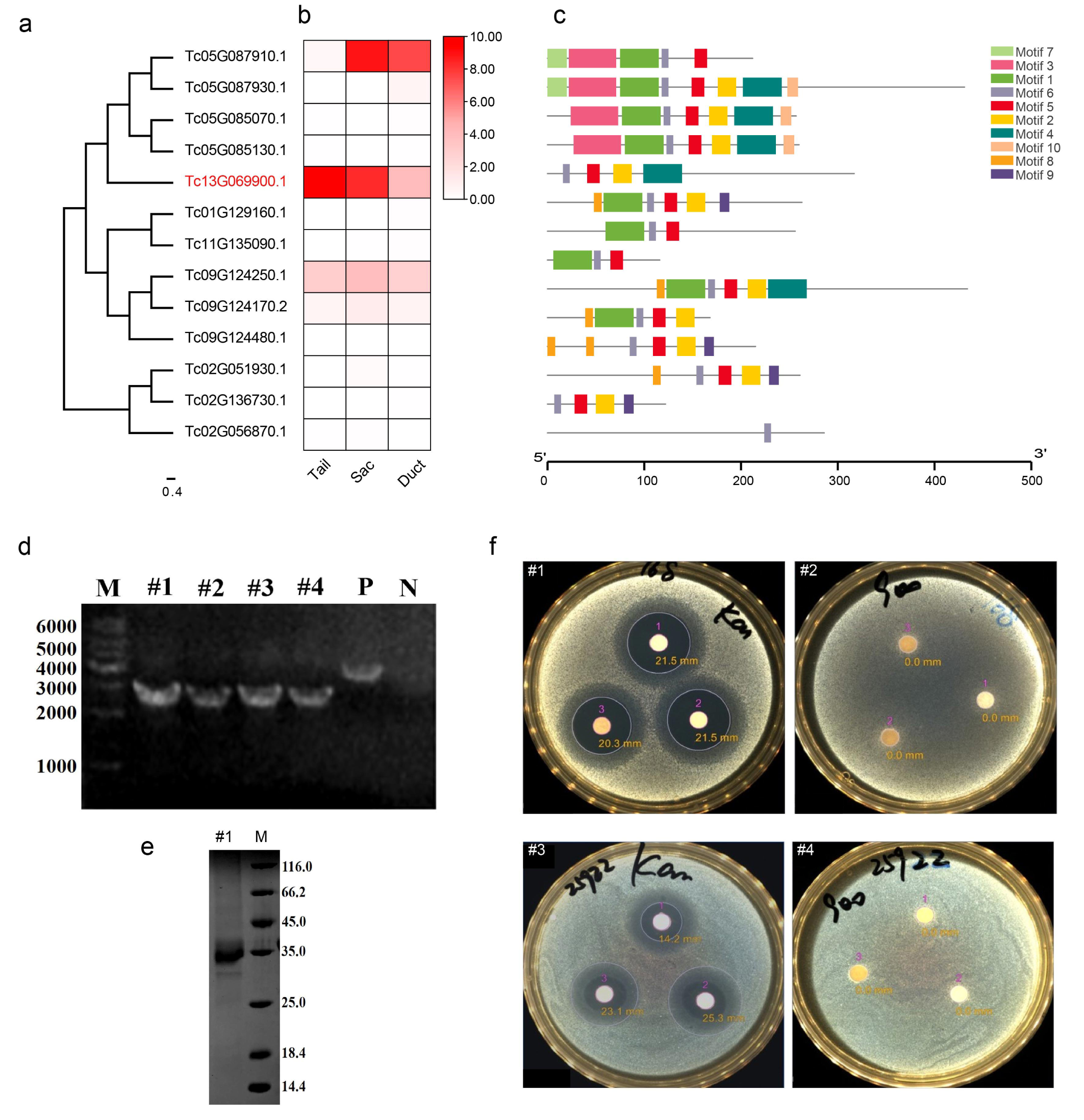
Venom protein function test. **a**, Phylogenetic tree of the CAP gene family to which Venom belongs. **b**, Expression heatmap of 13 CAP genes in the Tail, Sac and Duct. **c**, Conservative motif distribution. **d**, Agarose gel electrophoresis of the PCR product of the recombinant Bacmid plasmid. **e**, SDS–PAGE. **f**, The inhibition zone test by the disk diffusion test. #1: Bacillus subtilis 168_Kanamycin; #2: Bacillus subtilis 168_Venom; #3: Escherichia coli ATCC 25922_Kanamycin; #4: Escherichia coli ATCC 25922_Venom. The results indicated that Venom had no bacteriostatic function against these two bacteria.

**Supplementary Figure 15.**
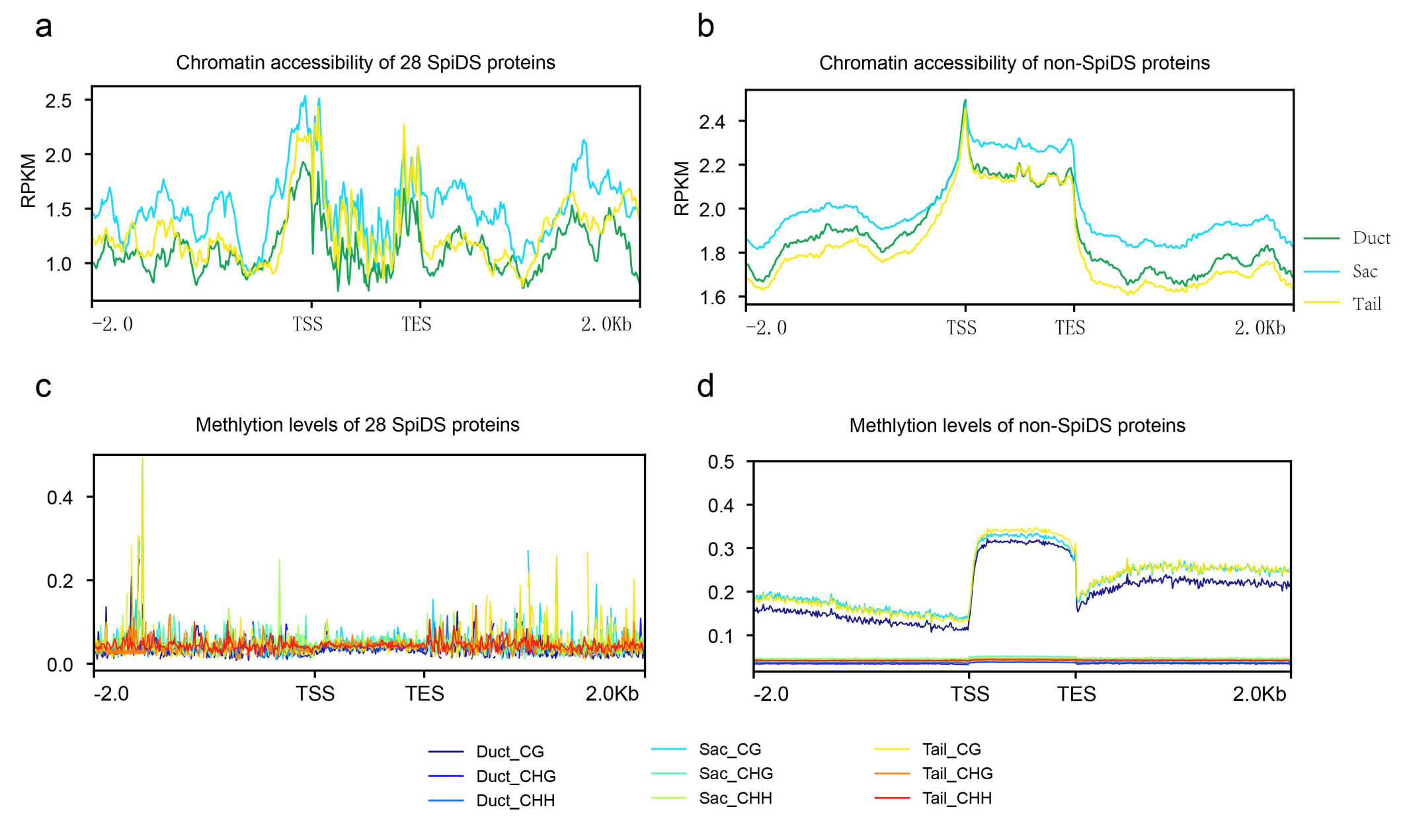
Chromatin accessibility and methylation levels for 28 spider dragline silk (SpiDS) and non-SpiDS genes.

**Supplementary Figure 16.**
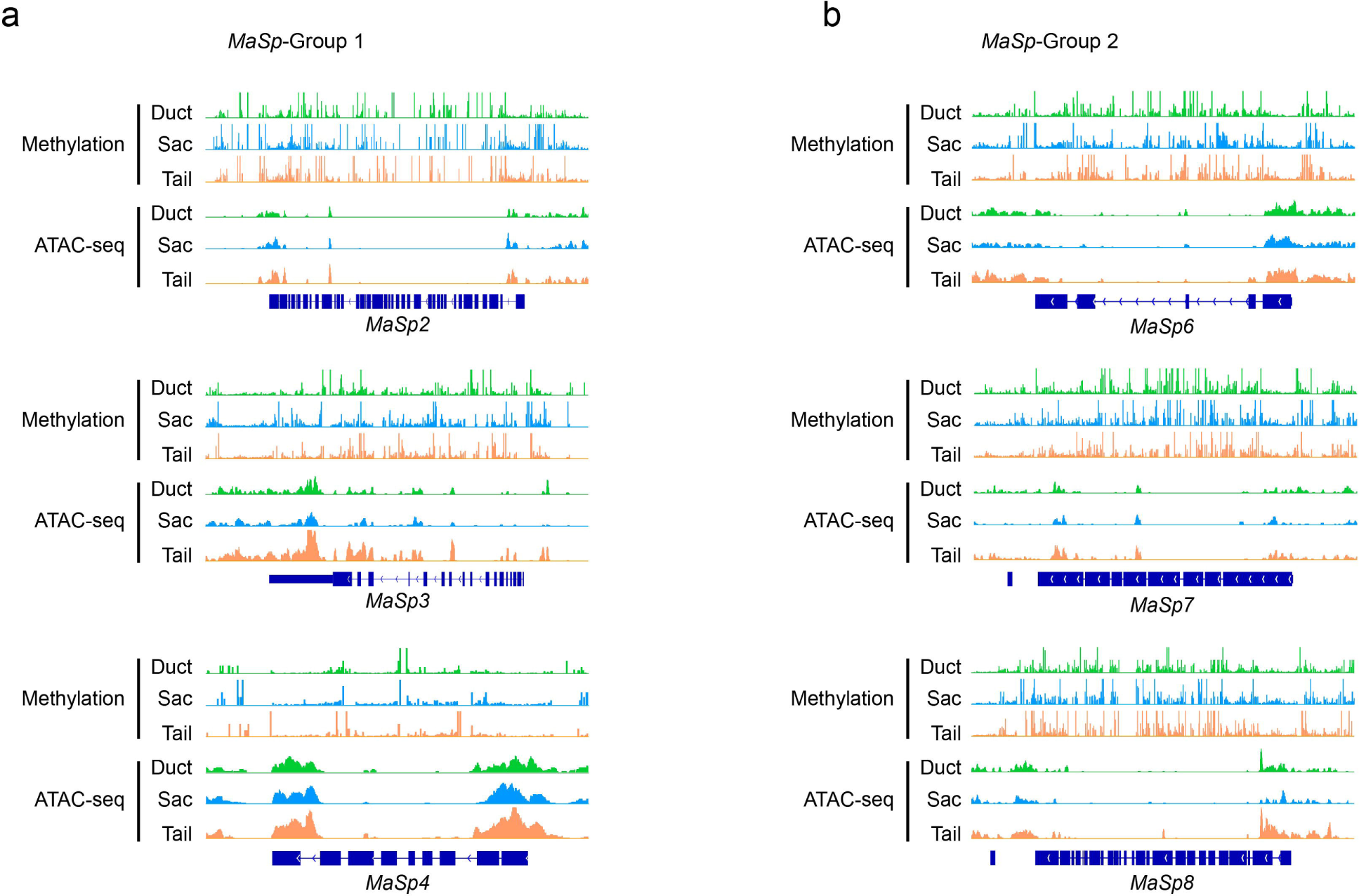
The methylation and chromatin accessibility peaks of *MaSp*-Group 1 (*MaSp2–4*) and *MaSp*-Group 2 (*MaSp6–8*).

**Supplementary Figure 17.**
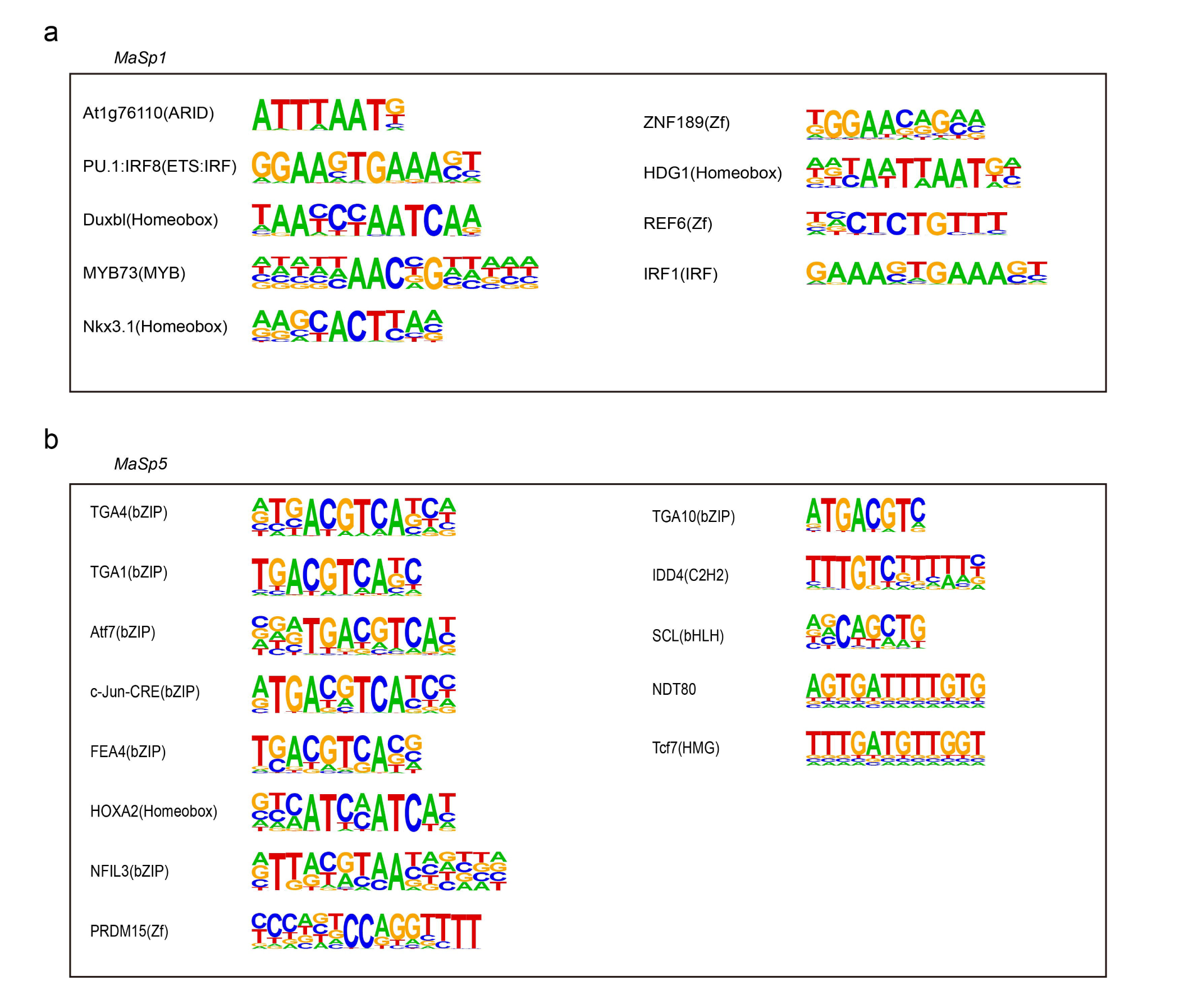
Specific motifs for *MaSp*-Group 1 and *MaSp*-Group 2. **a**, The specific motifs of *MaSp*-Group 1. **b**, The specific motifs of *MaSp*-Group 2. E-value < 1e^-10^.

**Supplementary Figure 18.**
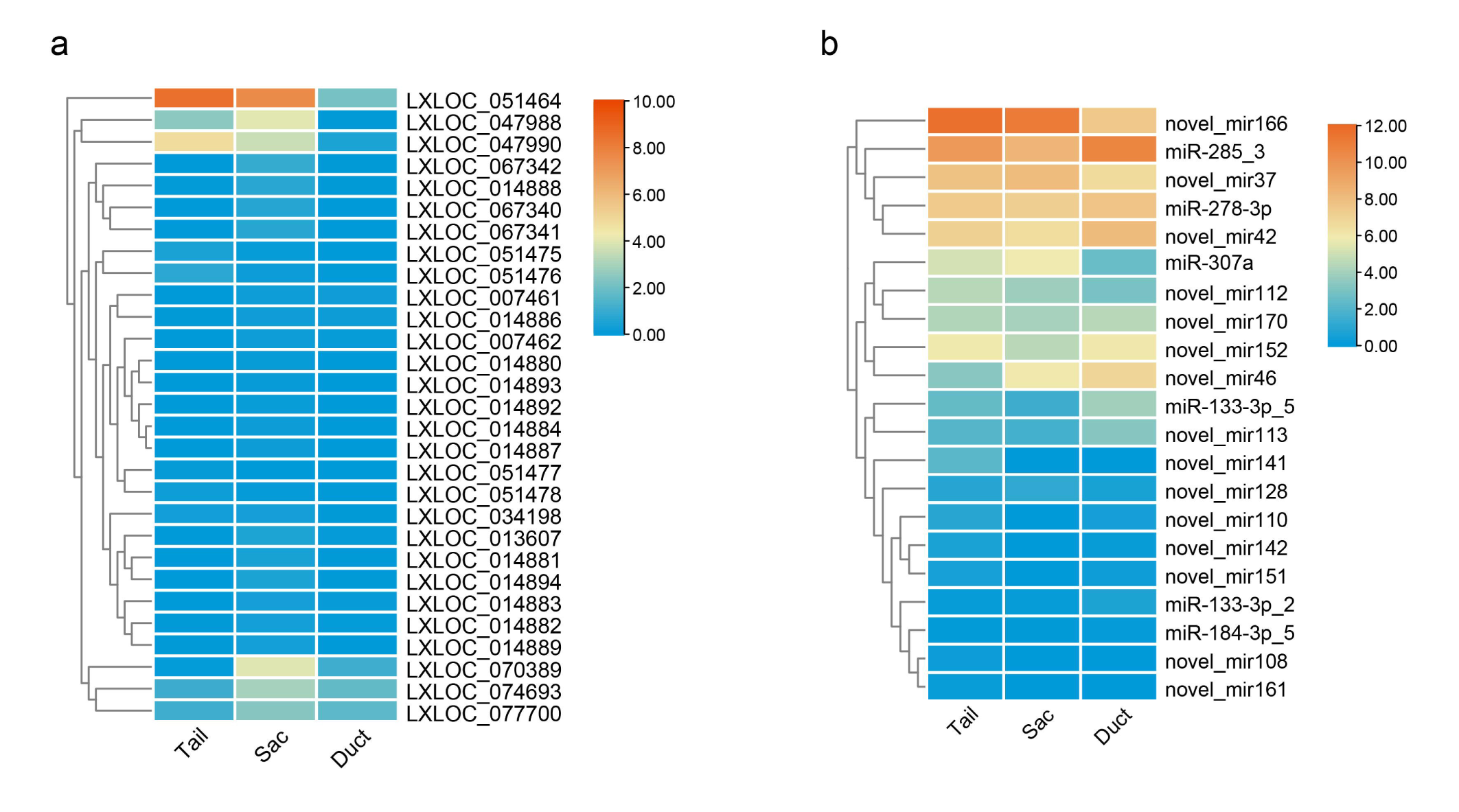
Expression heatmap of lncRNAs and miRNAs interacting with 28 SpiDS proteins. **a**, Expression heatmap of lncRNAs in the Tail, Sac, and Duct. **b**, Expression heatmap of miRNAs in the Tail, Sac, and Duct.

**Supplementary Figure 19.**
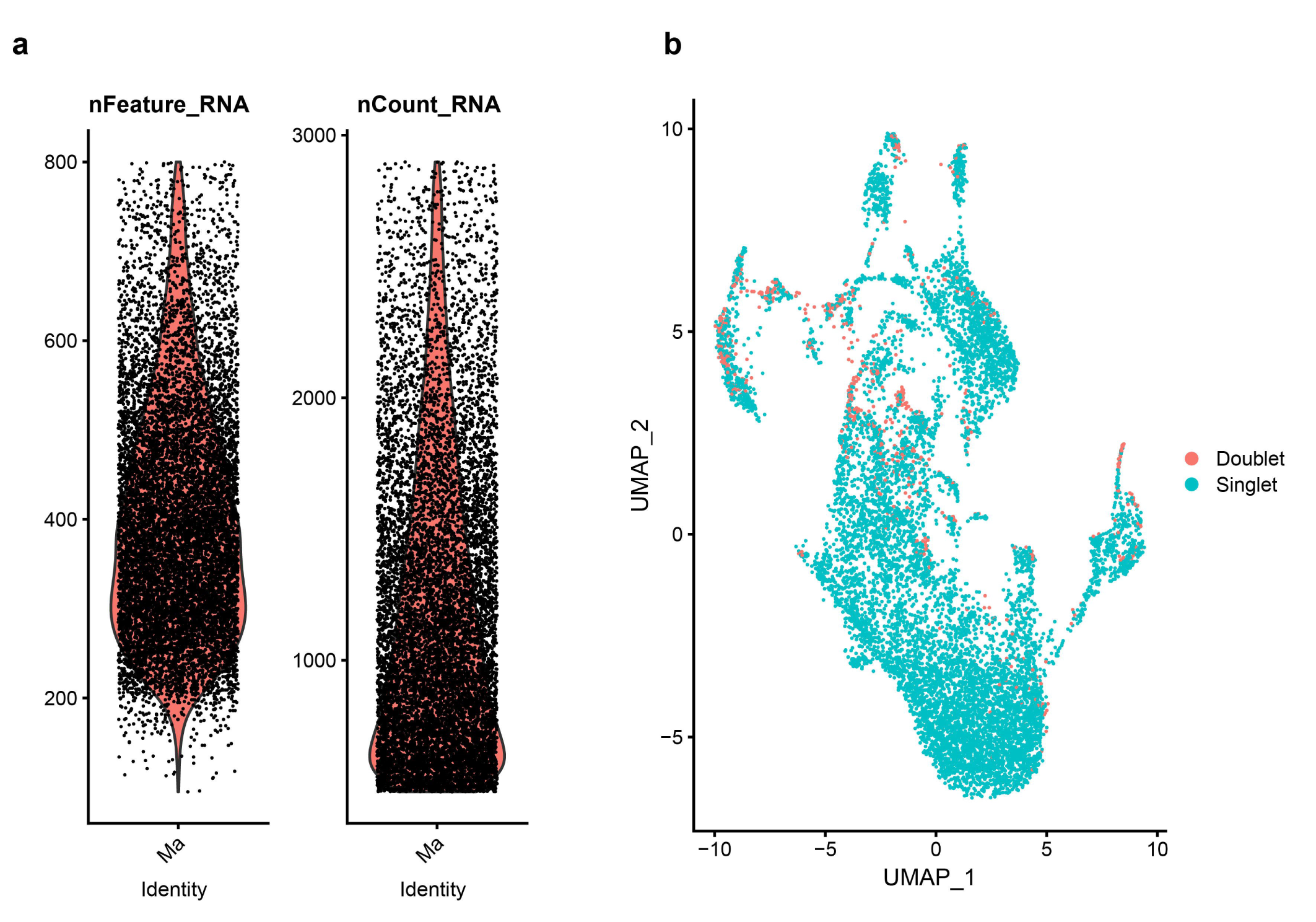
Ma scRNA-seq quality control. **a**, The number of gene expressions (left) and the number of UMIs (right) in each cell after quality control. **b**, Doublet cells were removed, and red dots represent possible doublet cells.

**Supplementary Figure 20.**
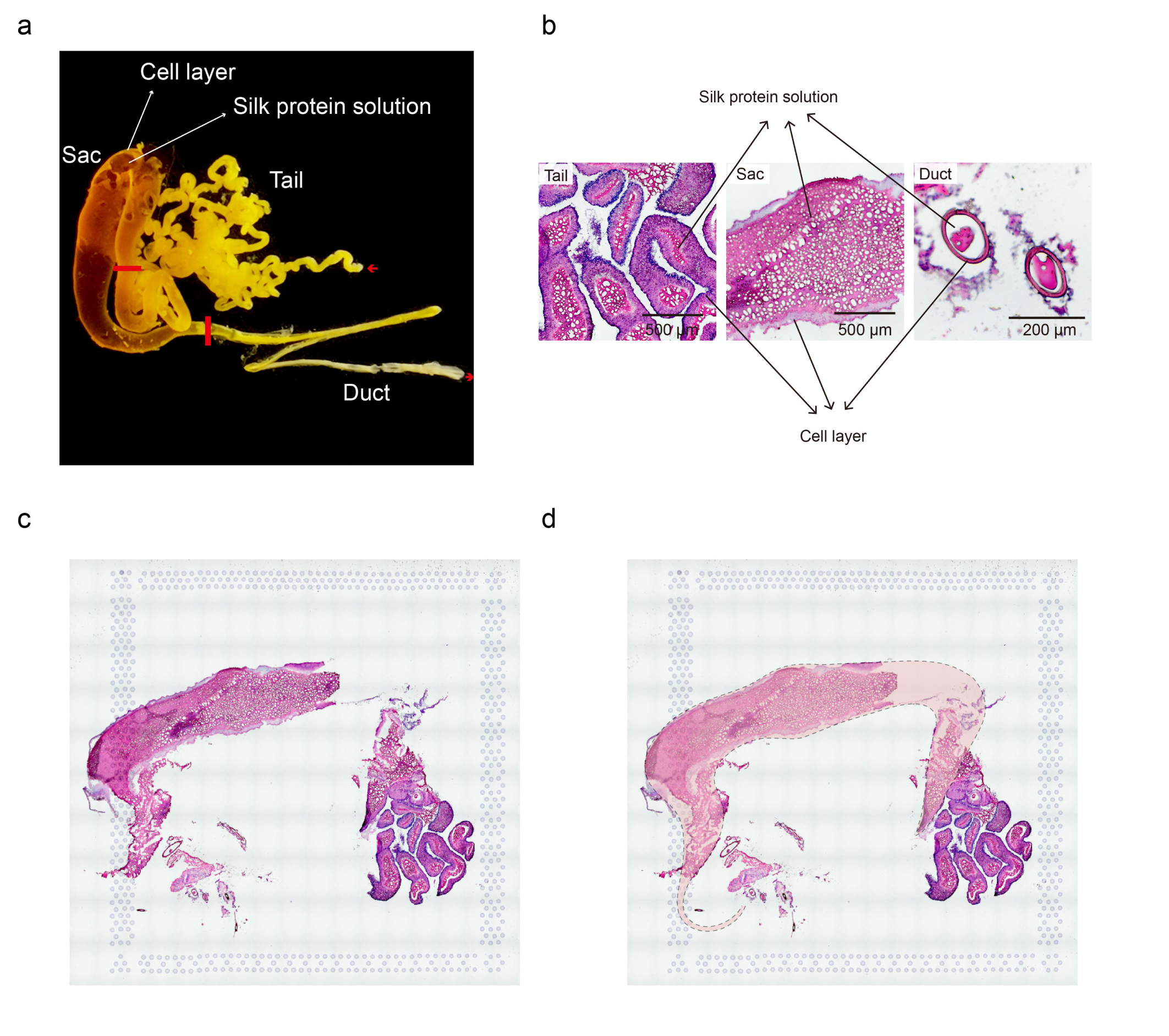
Morphological features and H.E. staining of Ma gland. **a**, Macroscopic appearance of the *T. clavata* Ma gland. **b**, Detail diagrams of MA silk gland by hematoxylin and eosin (H.E.) stained. Tail (longitudinal section), Sac (transverse section), and Duct (longitudinal section). **c**-**d**, H.E. staining of tissue sections for ST analysis. Dotted lines depict the outline of the Ma gland.

**Supplementary Figure 21.**
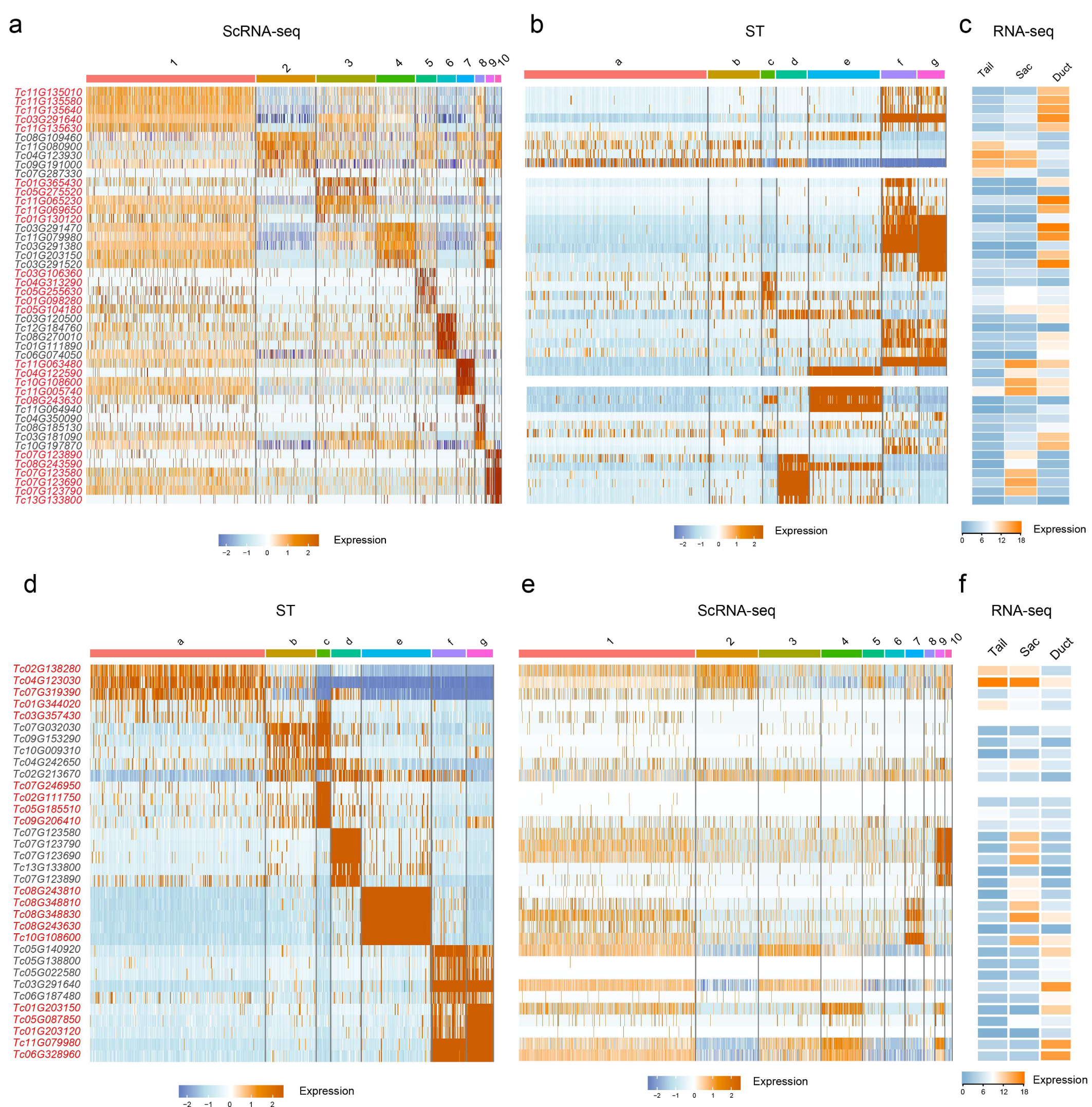
Expression heatmaps of marker genes. **a**-**c**, Expression heatmap of the top 5 SC marker genes in scRNA-seq, ST, and bulk RNA-seq. **d**-**f**, Expression heatmap of the top 5 ST marker genes in ST, scRNA-seq, and bulk RNA-seq.

**Supplementary Figure 22.**
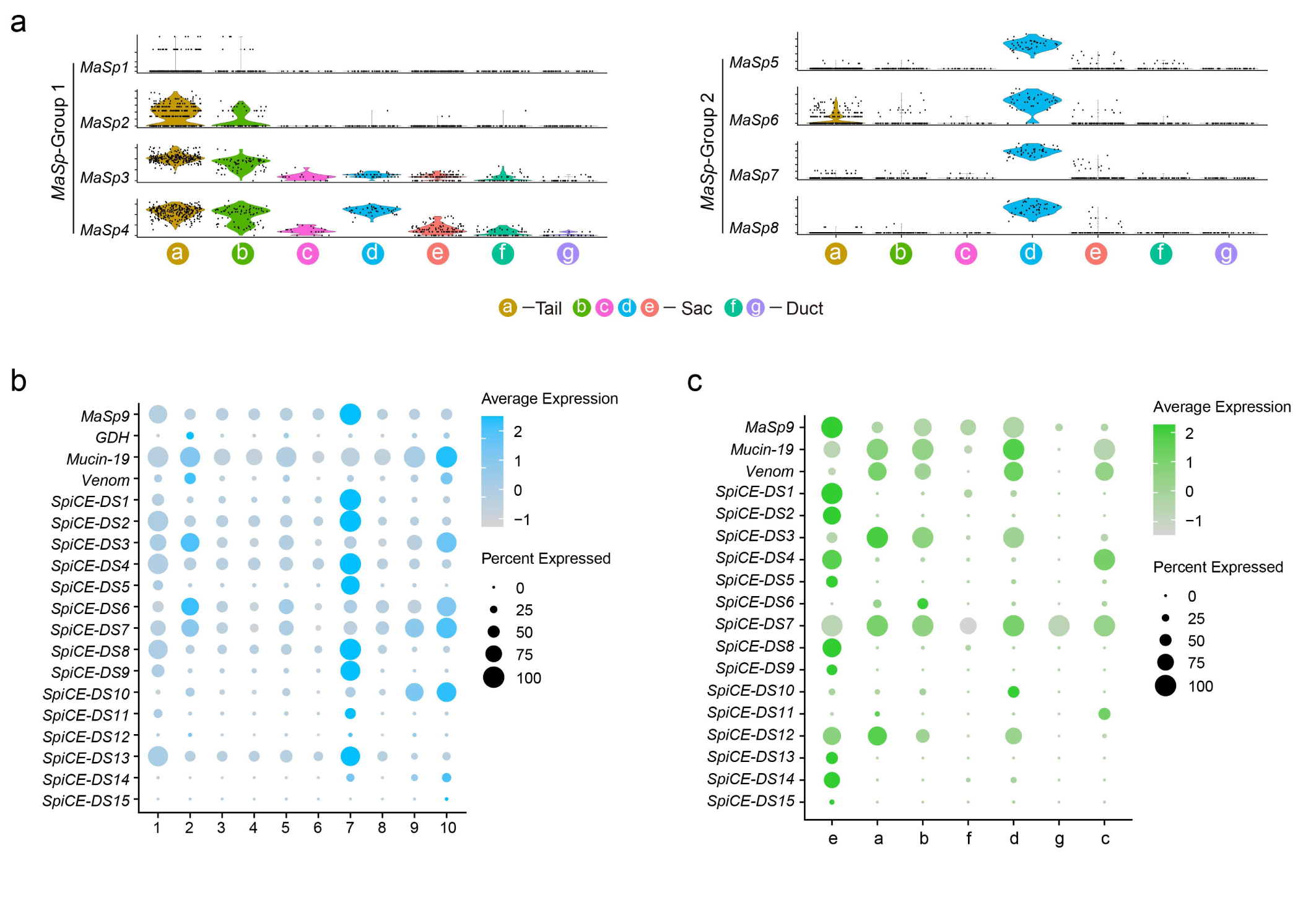
Expression patterns of SpiDS genes in scRNA-seq and ST. **a**, Violin plots of *MaSp* gene expression across ST clusters. **b**, Expression bubble plots of SpiDS genes in scRNA-Seq (*MiSp* was not expressed). **c**, Expression bubble plots of SpiDS proteins in ST (*MiSp* and *GDH* were not expressed).

**Supplementary Figure 23.**
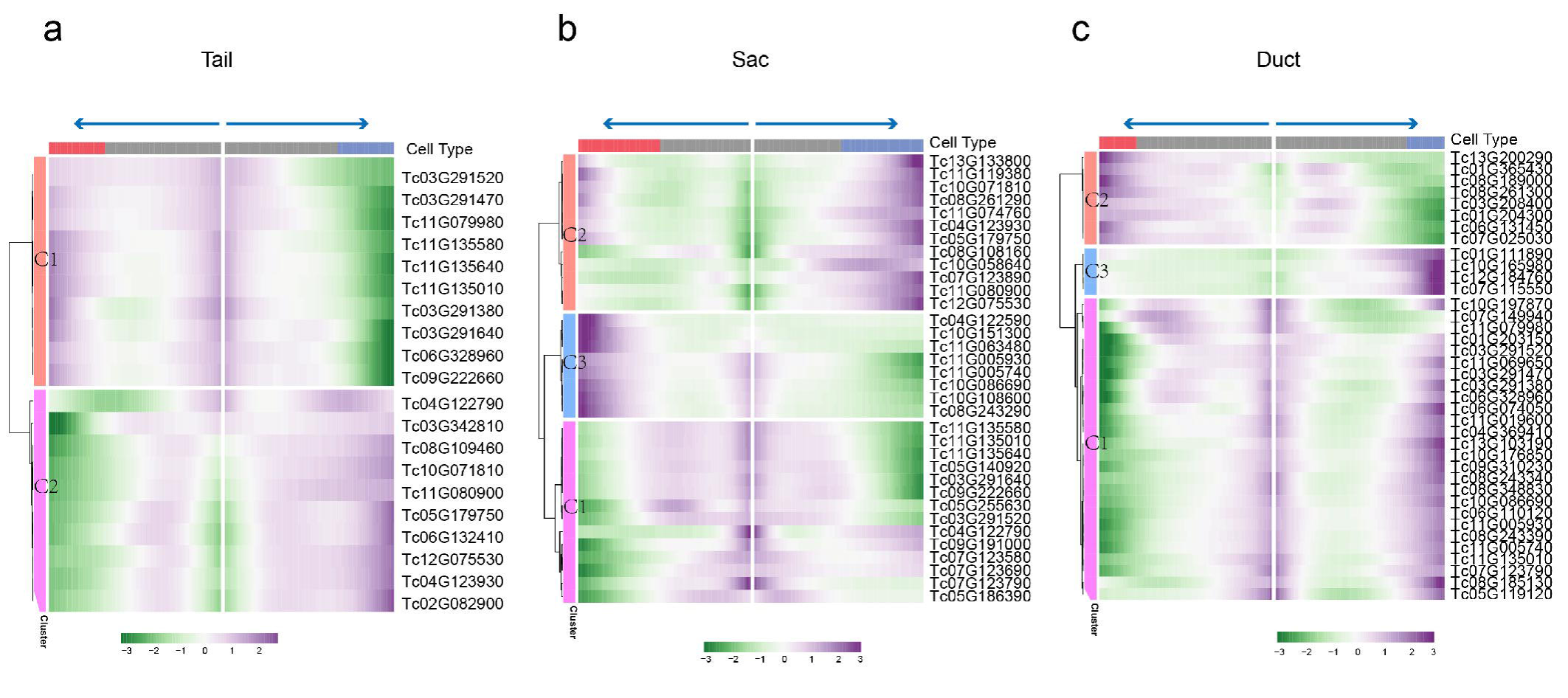
Developmental trajectories. **a**-**c**, Heatmap showing the expression of branch-dependent genes over pseudotime in the Tail, Sac and Duct. Representative marker genes are shown on the right of the heatmap. Both sides of the heatmap are the end of pseudotime.

**Supplementary Figure 24.**
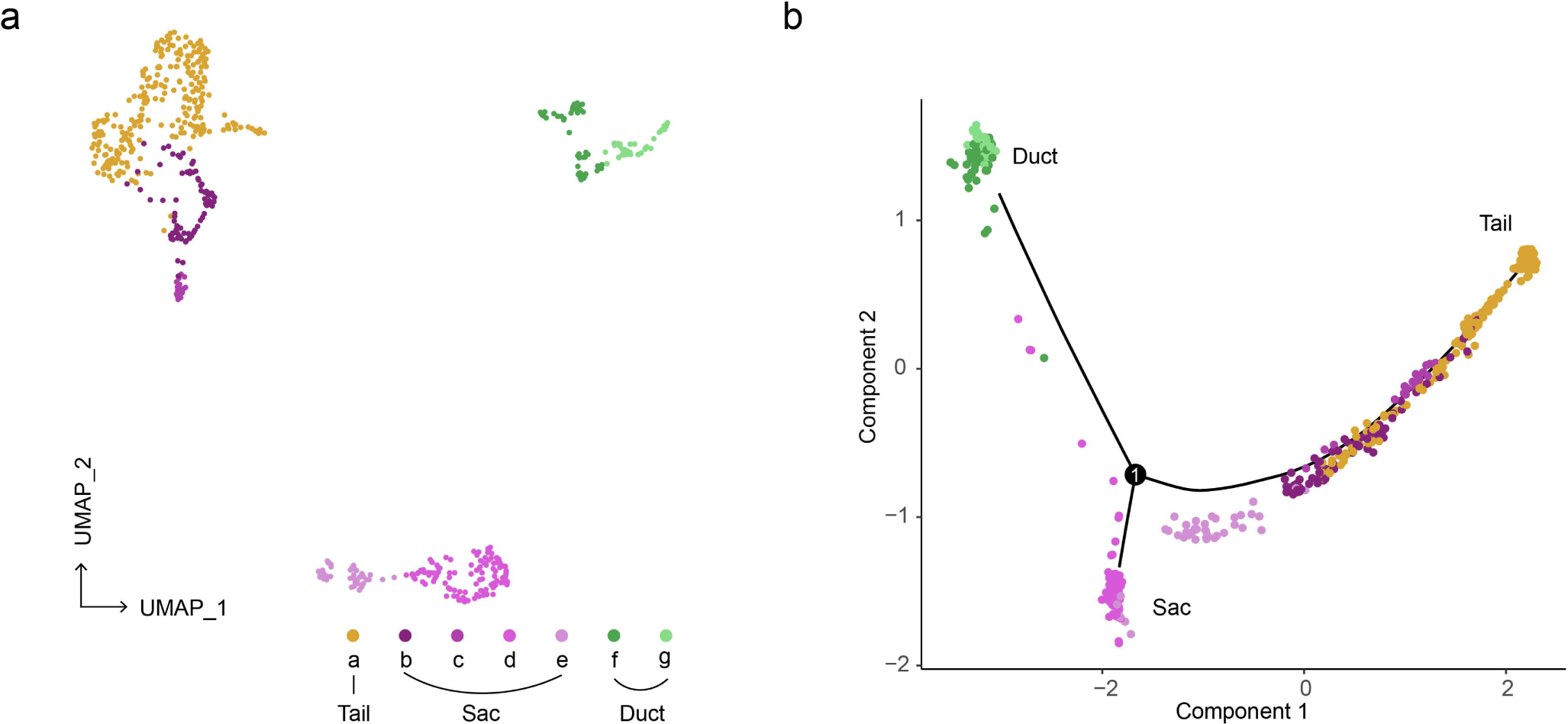
Spot clustering and developmental trajectory in the ST. **a**, UMAP visualization of seven putative clusters derived from 597 spots. **b**, Developmental trajectory of seven ST clusters.

**Supplementary Figure 25.**
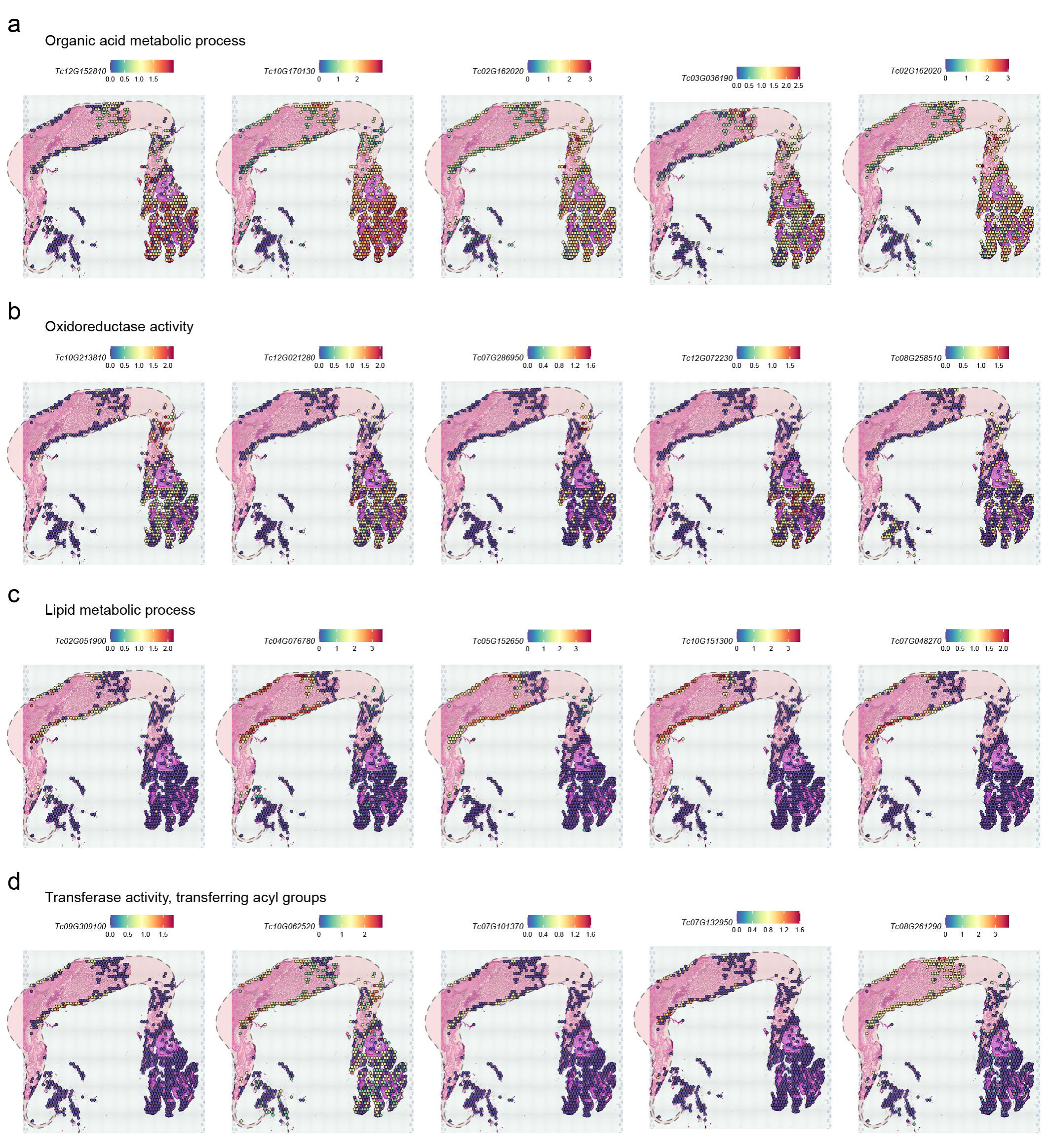
Feature plots of putative genes associated with dragline silk generation for Tail and Sac in ST. Tail: **a**, Organic acid metabolic process and **b**, Oxidoreductase activity. Sac: **c**, Lipid metabolic process and **d**, Transferase activity, transferring acyl groups.

**Supplementary Figure 26.**
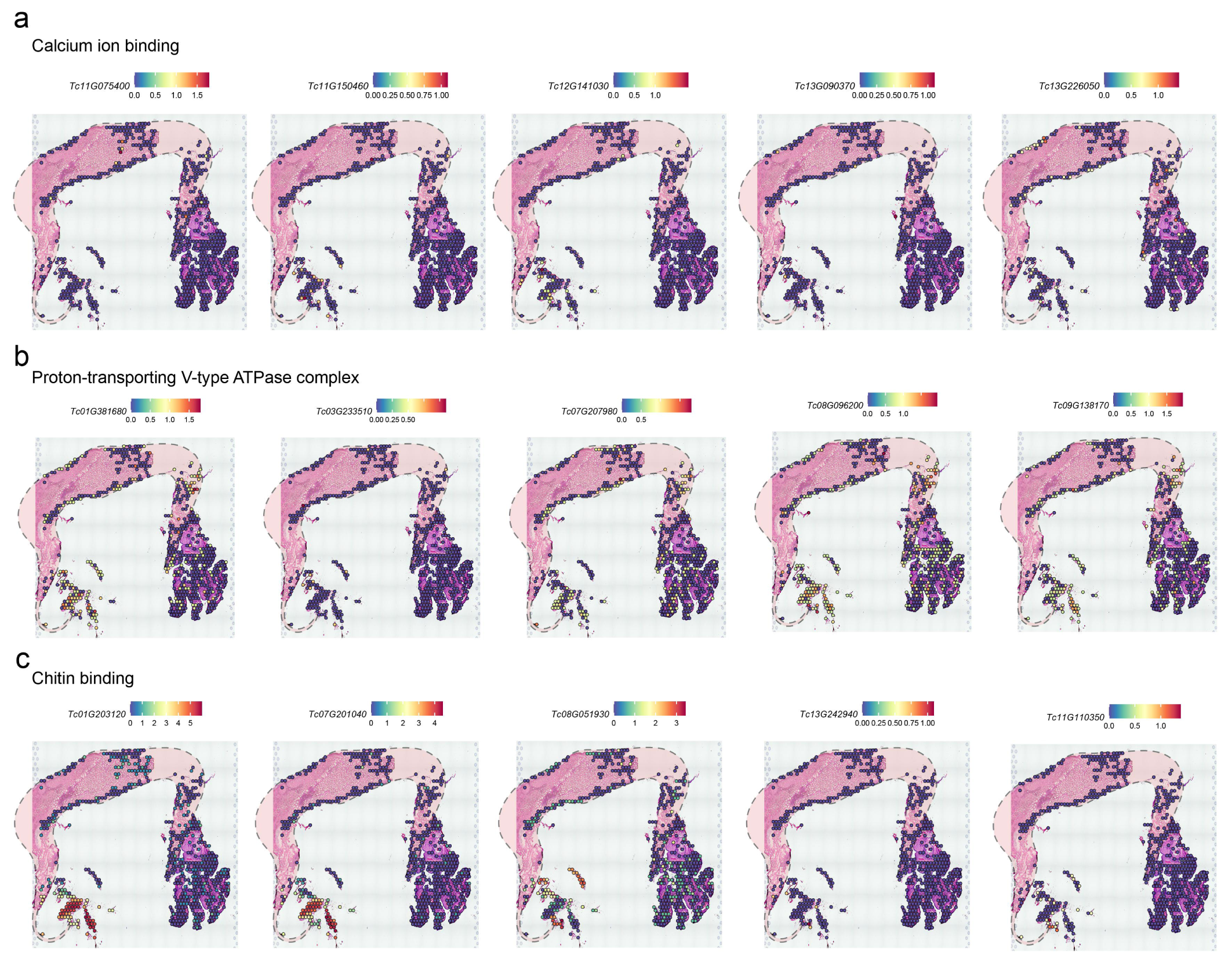
Feature plots of putative genes associated with dragline silk generation for Duct in ST. Duct: **a**, Calcium ion binding **b**, Proton-transporting V-type ATPase complex and **c**, Chitin binding.

**Supplementary Figure 27.**
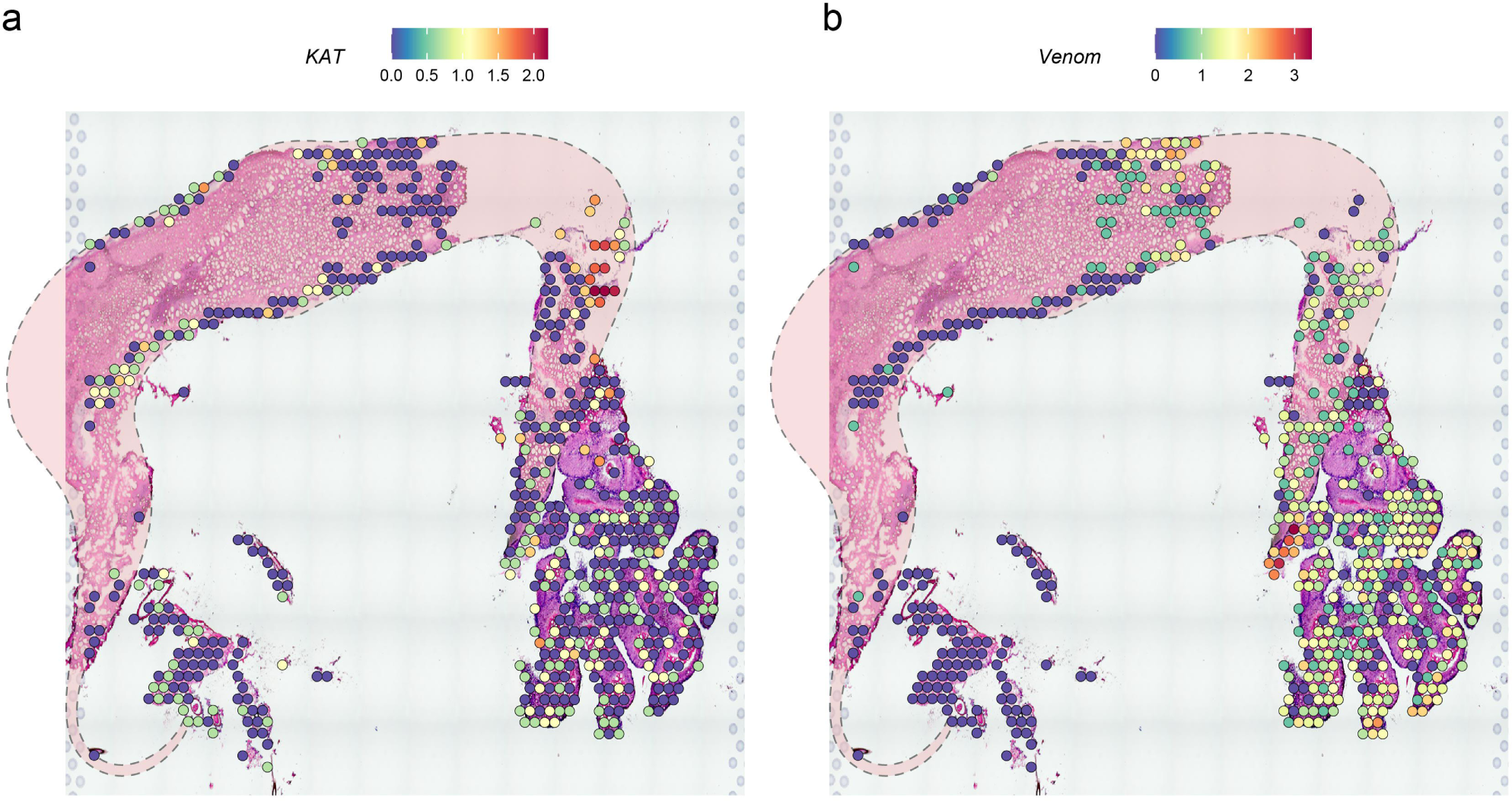
Feature plots of *KAT* and *Venom* in ST.

**Supplementary Figure 28.**
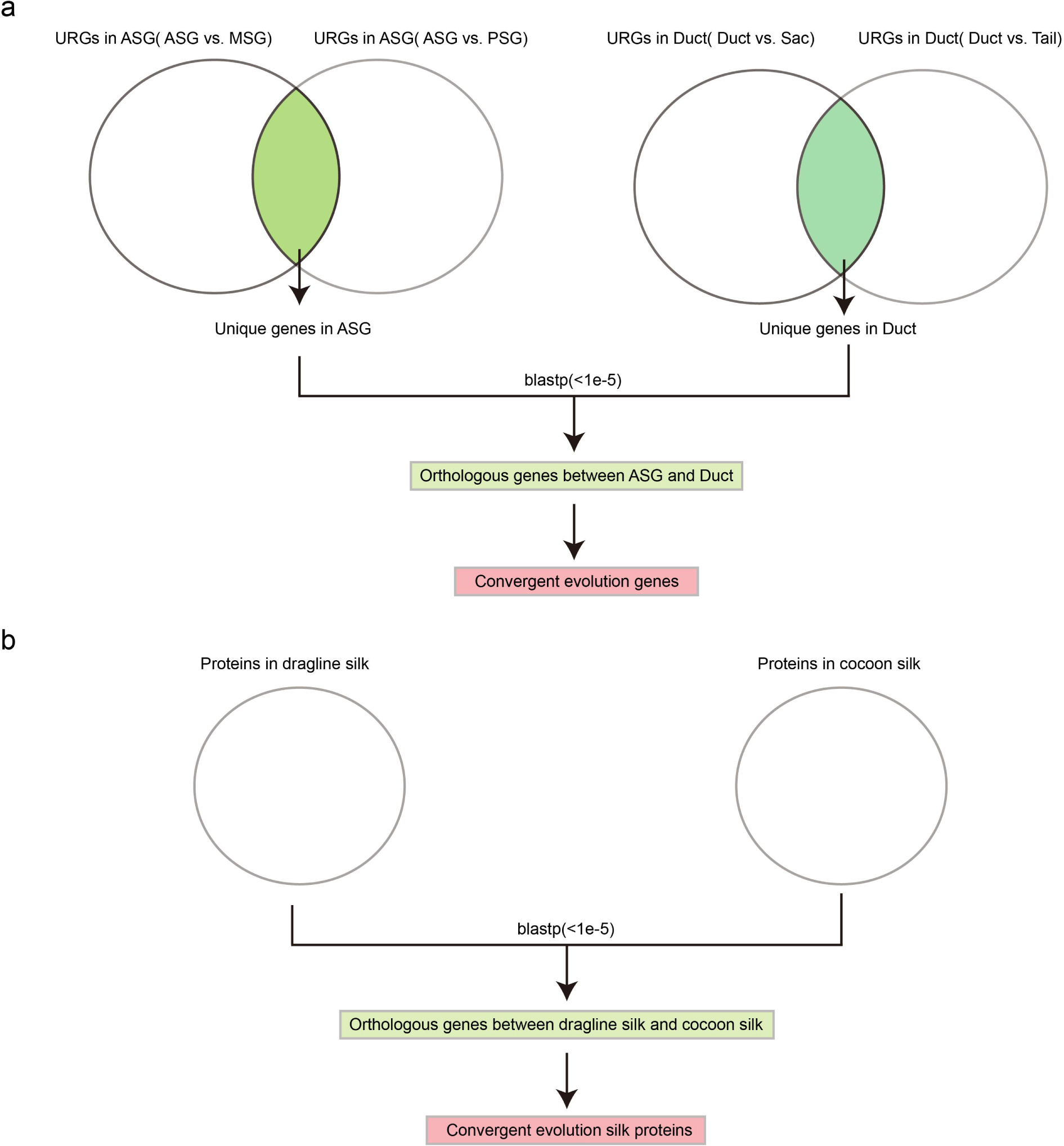
The pipeline of convergent evolution analysis. **a**, Flowchart of the screening of convergent genes in the silk gland for *B. mori* (ASG) and *T. clavata* (Duct). The same methods were used for MSG vs. Sac and PSG vs. Tail. **b**, Flowchart of screening convergent silk proteins for dragline silk (*T. clavata*) and cocoon silk (*B. mori*).

**Supplementary Figure 29.**
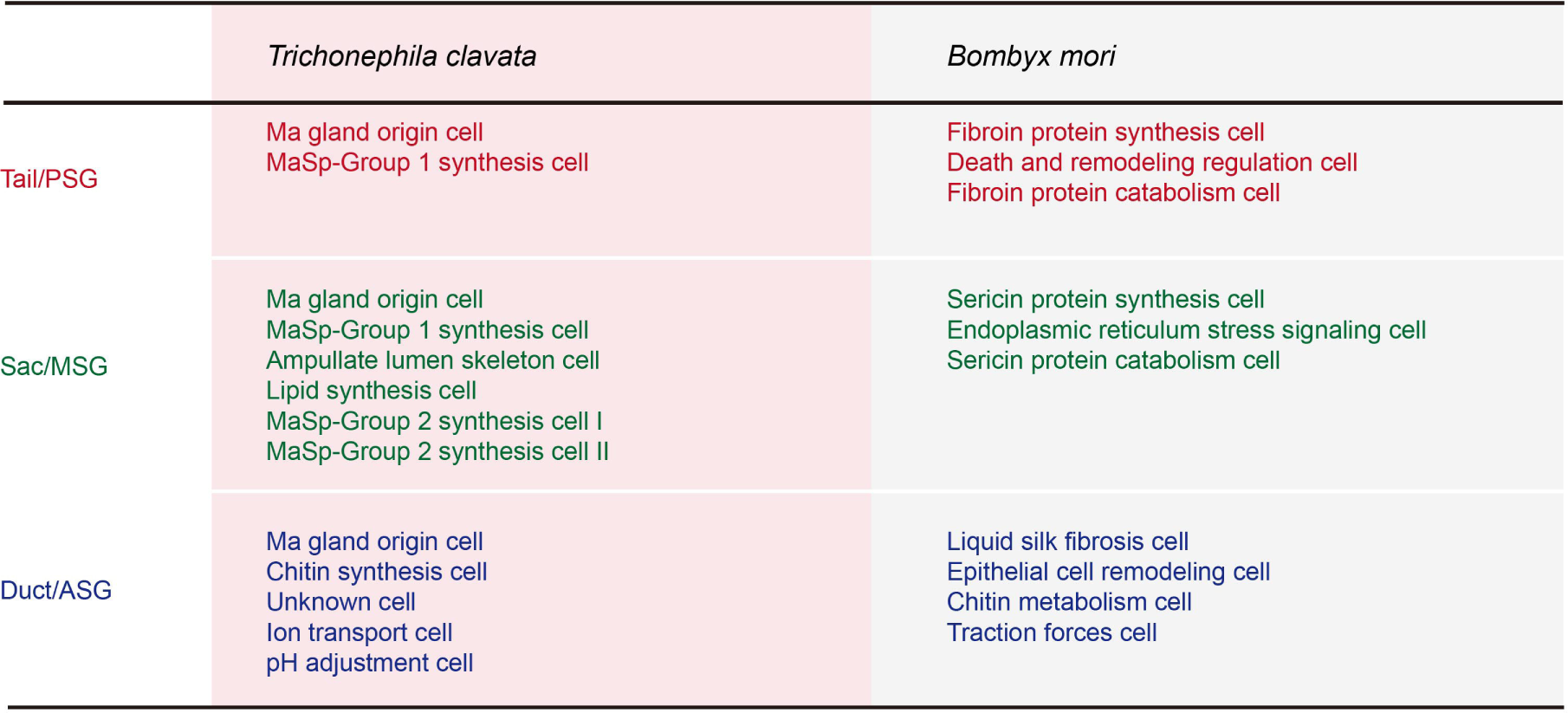
Comparison of single cell types of spider (*T. clavata*) Ma gland and silkworm (*B. mori*) silk gland. There was a similar number of silk gland cell types between *T. clavata* and *B. mori*, but differentiated annotations except for the chitin-related process in the Duct/ASG^58^.

**Supplementary Figure 30.**
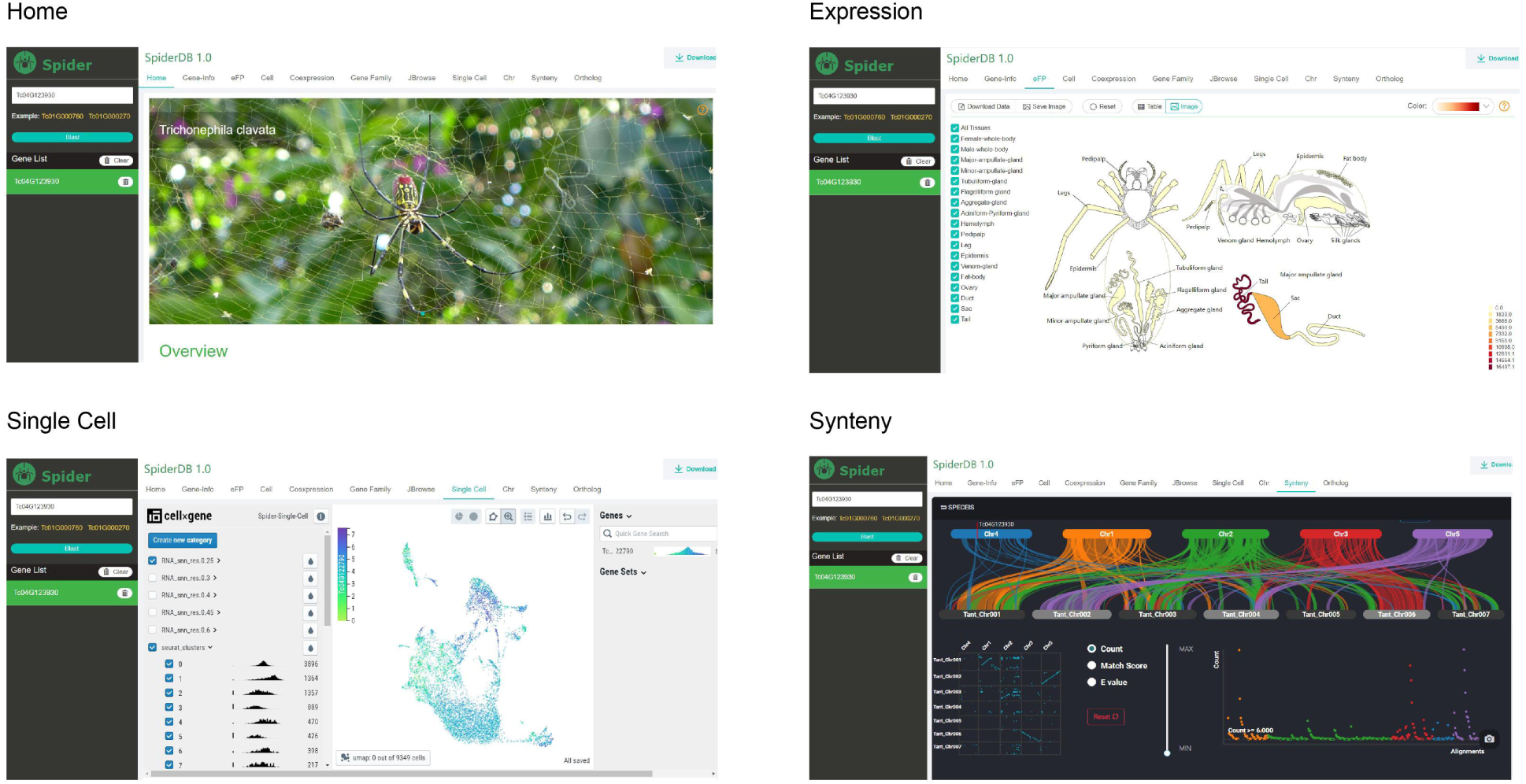
Several functional modules of SpiderDB. Only home page, expression, single cell, synteny modules were listed, for more function modules refer to spiderDB website (https://spider.bioinfotoolkits.net).

## Supplemental Tables

**All tables are in one excel file. Description:**

**Supplementary Table 1. Sample and sequencing Information**

**Supplementary Table 2. Summary statistics for genome survey**

**Supplementary Table 3. Summary statistics for T.clavata genome assembly**

**Supplementary Table 4. Summary statistics for Transposon Elements**

**Supplementary Table 5. Genome data resources for 12 species**

**Supplementary Table 6. Homologous evidence for identification of spider spdroin**

**Supplementary Table 7. N- and C-terminal sequence for identification of spider spdroin**

**Supplementary Table 8. Motif classes statistics for 28 spdroins in T.clavata**

**Supplementary Table 9. Statistics informations of strength, extensibility and stickiness related motifs**

**Supplementary Table 10. Silk protein abundance and Ma gland expression levels of 28 silk genes**

**Supplementary Table 11. The FPKM expression ratio of 28 SpiDS genes in Tail, Sac and Duct**

**Supplementary Table 12. Metabolite components of dragline silk**

**Supplementary Table 13. Metabolite component categories in the dragline silk**

**Supplementary Table 14. The GO enrichment results of the Duct, sac, and Tail-specific expression genes in transcriptome**

**Supplementary Table 15. Specific motifs in the upstream or downstream 2 kb of MaSp1 and MaSp5**

**Supplementary Table 16. Different types methylation ratios in different regions of the genome**

**Supplementary Table 17. Marker genes in single-cell clusters**

**Supplementary Table 18. Marker genes in ST clusters**

**Supplementary Table 19. The top5 GO annotations for each SC cluster**

**Supplementary Table 20. Convergent evolution of genes between silkworm silk gland and spider Ma gland**

**Supplementary Table 21. Silk protein abundance and silk gland tissue expression levels of 64 silk genes in silkworm**

**Supplementary Table 22. Metabolite components of cocoon silk**

